# The black-legged tick *Ixodes scapularis* detects CO_2_ without the Haller’s organ

**DOI:** 10.1101/2023.08.03.551779

**Authors:** Carola Städele

## Abstract

Both male and female ticks have a strong innate drive to find and blood-feed on hosts. Carbon dioxide (CO_2_) is considered a critical behavioral activator and attractant for ticks and an essential sensory cue to find hosts. Yet, how CO_2_ activates and promotes host-seeking in ticks is poorly understood. We studied CO_2_ responses in the black-legged tick *Ixodes scapularis*, the primary vector for Lyme disease in North America. Adult males and females were exposed to 1, 2, 4, or 8% CO_2_, and changes in walking behavior and foreleg movement were analyzed. We find that CO_2_ is a potent stimulant for adult *Ixodes scapularis*, even at lower concentrations (1%). Behavioral reactions depend on the animal’s state: Walking ticks increase their walking speed, while stationary ticks start to wave their forelegs and begin to quest – both behaviors resembling aspects of host-seeking. Furthermore, *Ixodes scapularis* has no clear concentration preference and is not tuned more robust to breath-like CO_2_ concentrations (∼4%) than to the other concentrations tested. As soon as the CO_2_ level is above a certain threshold, *Ixodes scapularis* react, indicating that CO_2_ acts as a behavioral activator and can be used as a long-distance cue to detect approaching hosts. Moreover, we provide convincing evidence that the foreleg Haller’s organ is not necessary for CO_2_ detection. Even with disabled or amputated Haller’s organ, *Ixodes scapularis* respond robustly to CO_2_, signifying that there must be CO_2_-sensitive structures important for tick host-seeking that have not yet been identified.

## INTRODUCTION

The survival and maturation of ticks require finding and blood-feeding on hosts. Both male and female ticks need at least one blood meal in each of their three development stages (larva, nymph, adult) to become sexually mature and reproduce (Apanaskevich et al., 2014). Especially in lay literature, the approximately 4% CO_2_ of mammalian breath is often cited as a critical behavioral activator and attractant for ticks and an essential sensory cue for host-seeking. However, few tick species have been tested for their CO_2_ sensitivity, and quantitative laboratory studies are lacking.

CO_2_ is a kairomone used by many hematophagous parasites to sense the proximity of a potential host (summarized by Chaisson and Hallem, 2012). For instance, CO_2_ exposure elicits a range of contextual effects in various mosquito species that are thought to promote host-seeking behavior. These effects include activation of flight (Eiras and Jepson, 1991), increased attraction to body heat (Liu and Vosshall, 2019; McMeniman et al., 2014), and enhanced steering responses to vertically elongated dark visual objects (Vinauger et al., 2019). Whether CO_2_ activates and promotes host-seeking in ticks has not been well studied. However, there is evidence that exposure to CO_2_ elicits behavioral responses related to host-seeking.

The two dominant host-seeking strategies for ticks are active hunting and passive ambushing (Apanaskevich et al., 2014). Hunting ticks actively invade the habitat of hosts while ambushing ticks quest by climbing up the vegetation to wait for passing hosts, typically with extended forelegs. Many economically important tick species, including the black-legged tick *Ixodes scapularis* studied here, predominantly quest for hosts. It is well known among tick field ecologists that CO_2_ traps efficiently capture tick species that hunt for hosts (Garcia, 1969, 1962; Guedes et al., 2012; Guglielmone et al., 1985; Norval et al., 1987; Oliveira et al., 2000). For example, the lone star tick Amblyomma americanum, known for its aggressive hunting behavior, is attracted by CO_2_ traps from up to 21 meters away (Wilson et al., 1972). However, species that predominantly quest to find hosts are only sporadically captured with CO_2_ traps (Garcia, 1962; Guedes et al., 2012).

With about 850 species of ticks living in habitats from the cold Arctic to arid deserts, it is not surprising that anatomy, seasonal activity patterns, host preference, and host-seeking strategies vary across species. For example, the winter tick, *Dermacentor albipictus*, is a so-called one-host tick that spends its life almost entirely on the same host (Howell, 1940). In contrast, *Ixodes scapularis* is a three-host tick that feeds on a different host during each life stage and leaves the host to molt (Keirans et al., 1996; Soneshine and Roe, 2014). Larvae preferably feed on small mammals such as rodents, while nymphs and adults prefer larger hosts such as deer, dogs, horses, and humans (Keirans et al., 1996). Due to this three-host lifestyle, *Ixodes scapularis* efficiently spreads blood-borne diseases such as Lyme borreliosis (Eisen and Eisen, 2018). How CO_2_ contributes to host-seeking or selection in this notorious disease vector is not understood.

Most studies examining CO_2_ attractiveness for ticks have not aimed to explain the physiology of CO_2_ detection. Instead, many just tested the effectiveness of CO_2_ as bait and focused on quantifying changes in locomotion, i.e., counted how many ticks actively walked towards a trap (Garcia, 1969, 1965, 1962; Gray, 1985; Guedes et al., 2012; McMahon and Guerin, 2002; Miles, 1968; Nevill, 1964; Norval et al., 1987; Wilson et al., 1972). Since tick species that predominantly quest for hosts are rarely captured with CO_2_ traps, it is reasonable to ask whether the presence of CO_2_ motivates questing ticks to walk and whether quantifying changes in locomotion is sufficient to describe CO_2_-induced behavioral responses. Van Duijvendijk and colleagues (van Duijvendijk et al., 2017), for example, tested *Ixodes ricinus* nymphs, the European cousin of *Ixodes scapularis*, for CO_2_ attractiveness using a Y-maze choice olfactometer. Attractiveness was assessed by counting the number of nymphs that actively approached the odor arms. Possible non-walking-related behavioral reactions in response to CO_2_ were not evaluated. Less than 25% of the animals tested walked during the selection test, and the few walking animals did not prefer CO_2_ over air. The sparse number of walking ticks suggests that CO_2_ might not induce walking in *Ixodes ricinus* nymphs. Questing larvae of *Dermacentor albipictus*, for instance, do not start to walk when exposed to CO_2_ but remain in their questing position (Garcia, 1969).

Our approach differs significantly from most previous studies in that it includes observations of behavioral reactions other than changes in locomotion. We demonstrate that CO_2_ is a potent stimulant for adult male and female *Ixodes scapularis* and that behavioral responses depend on the state of the animal. Walking animals tend to change their walking speed, while stationary animals remain stationary but start to wave their forelegs. This latter behavior was striking and resembled the described foreleg waving of questing ticks, assumed to support CO_2_ detection. The forelegs of ticks carry a specialized multisensory structure, the Haller’s organ (HO), which has been implicated in CO_2_ detection (Carr et al., 2017; Steullet and Guerin, 1992). However, we provide strong evidence that the HO is not necessary for CO_2_ detection and that other CO_2_-sensitive structures, not yet identified, must be present on the tick body.

## RESULTS

### *Ixodes scapularis* responds robustly to CO_2_, but without a clear concentration preference

To determine behavioral responses and sensitivity to CO_2_, we exposed unfed adult *Ixodes scapularis* to different CO_2_ concentrations (1, 2, 4, and 8%). An air puff with the same duration as the 8% CO_2_ treatment was used as a negative control to distinguish behavioral responses to CO_2_ from changes in airflow. The treatment order was not randomized in these experiments. Air puff controls were always first, followed by ascending CO_2_ treatments. We tested 50 ticks, 25 males and females, in groups of five. In each experiment, five ticks were placed inside a Plexiglass arena (Figure 1A) and video monitored for 10 minutes (5 minutes in ambient air and during treatments).

**Figure 1:**
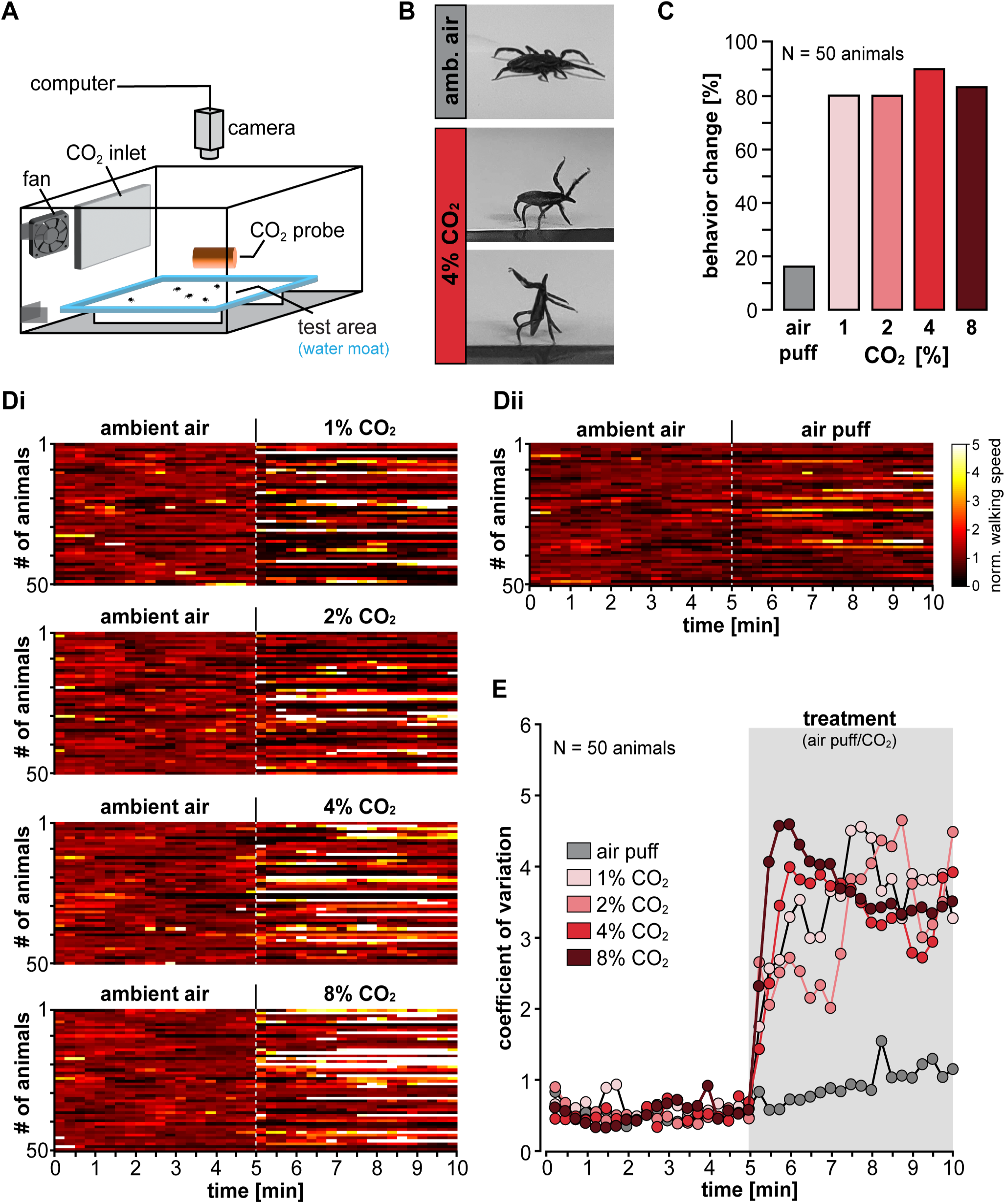
*Ixodes scapularis* respond robustly to CO_2_ at all concentrations tested. **(A)** Schematic of the multi-animal CO_2_ test arena. **(B)** Example postures of an adult female *Ixodes scapularis* before and during CO_2_ exposure. CO_2_ exposure can elicit questing (= foreleg raising or foreleg and body raising). **(C)** Analysis of behavioral changes of 50 adult *I. scapularis* (25 males and 25 females) during exposure to an air puff (=negative control) or CO_2_ (1, 2, 4, and 8%; shades of red). Data were obtained by manual video analysis 2 minutes before and after the onset of the treatments. **(D)** Analysis of changes in normalized walking speed before (ambient air) and during exposure to different CO_2_ concentrations/air puff (N = 50 animals). Brighter colors represent faster walking speeds. The dashed gray line depicts the beginning of the treatment (air puff or CO_2_). Data were obtained by calculating walking velocity from x/y positions (computer tracking) and do not contain potential changes in foreleg/body position. Same animals as in panel B. Bin width = 30 frames, sampling rate = 2 frames/s. Data from each animal were normalized to the respective mean of bins 1 to 10. **(E)** Coefficient of variation for the data presented in panel D. Dots represent mean values (50 animals). The gray rectangle illustrates the start and

The animals were allowed to behave freely, and accordingly, we observed different behavioral states in control conditions (ambient air). Some animals walked, while others were stationary and did not move. We took an unbiased approach and looked for changes in behavior for walking and non-walking animals alike.

The two most prominent behavioral reactions of *Ixodes scapularis* to CO_2_ were changes in walking speed and extensive foreleg waving, often associated with changes in posture. Foreleg waving was easily identifiable in non-walking animals, whose forelegs typically pointed forward and rested on the ground in ambient air (Figure 1B, top). In the presence of CO_2_, many of these animals began to raise and wave their forelegs (Figure 1B, middle). In some cases, animals would erect their entire body while waving with their forelegs (Figure 1B, bottom). During walking, the forelegs were often pointed forward, constantly moving, and touching the ground at irregular intervals. The animals were video-monitored from above. Due to the extensive foreleg movement during walking, we could not resolve alterations in foreleg position for walking animals in these experiments. It is likely, however, that CO_2_ elicited increased foreleg waving in walking animals, too.

To quantify how many animals responded to the treatments, we scored changes in walking speed and foreleg movements with either 0 (no change) or 1 (behavioral change) over 2 minutes in ambient air and CO_2_, respectively. Overall, the air puff did not strongly affect *Ixodes’* behavior. Only 16% of the animals responded (Figure 1C). However, *Ixodes scapularis* showed high sensitivity to all tested CO_2_ concentrations. 80% of the 50 animals showed behavioral responses to 1 and 2% CO_2_. Most behavioral changes were observed at 4% CO_2_ (90%) and slightly less at 8% (82%). Our data thus show that *Ixodes scapularis* has no clear concentration preference and is not tuned stronger to breath-like CO_2_ concentrations (∼4%) than to the other concentrations tested. As soon as the CO_2_ level was above a certain threshold, *Ixodes scapularis* reacted - supporting the hypothesis that CO_2_ acts as an activator.

To test for sex-specific responses, we analyzed males and females separately (Supp. Figure S1). We did not observe substantial differences. Males and females responded similarly to the tested CO_2_ concentrations (Supp. Figure S1), with males showing slightly more reactions at 1% (88% versus 72% females) and females at 4% CO_2_ (96% versus 84% males). We thus pooled male and female data in all further analyses. The sex ratio was 50% males and 50% females in all experiments.

*Ixodes*’ behavior was quite variable in the control condition (ambient air): Animals displayed various behaviors that changed with time. For example, the animals often did not walk continuously but paused at irregular intervals before resuming locomotion. Consequently, a simple comparison of walking speeds before and during treatment showed no difference due to the high variability (Supp. Figure S2A). However, we found sudden decreases in walking speed at the onset of the stimulus for all applied treatments, including the air puff (dark band in Supp. Figure S2A). Manual video analysis revealed that many ticks stopped briefly and began to wave with their forelegs. Since these initial speed decreases were short and lasted roughly 10 seconds, we refer to them as startle response. Because 30% of the ticks already showed startle responses during the control air puff (Supp. Figure S2B), we hypothesize that these were partly mechanosensory and elicited by changes in airflow.

Despite the high variability, we aimed to quantify behavioral changes in walking speed in more detail. To detect possible changes, we normalized each animal’s data to its average walking speed during the first 150 seconds of the experiment (10 bins). Thus, an increase in walking speed resulted in numbers greater than 1 and a decrease in numbers smaller than 1. Figure 1D shows the change in walking speed as heatmaps for the 50 animals tested throughout the 10-minute experiments. The number of animals that increased their walking speed rose with higher CO_2_ concentrations, evident from a shift towards lighter colors in the heatmaps. However, some animals also decreased their walking speed or stopped entirely in response to CO_2_ (darker colors in heatmaps). Note that reductions in walking speed are underrepresented in the heatmap color space as they can only range between 1 (initial walking speed) and 0 (animal is stationary). In contrast, increases in walking speeds ranged between 1 and 5 in our experiments.

To eliminate this imbalance, we calculated the coefficient of variation (CV) from the normalized data (Figure 1E). Any deviation from the initial walking speed will increase the CV, independently of whether the animal slowed down or walked faster. The control air puff caused no substantial effect, and CV progression over time was almost linear. In contrast, CVs for the tested CO_2_ concentrations increased rapidly at the CO_2_ onset, with most of them reaching their maximum about one minute after the start of the treatment. To conclude, our data show that *Ixodes scapularis* respond robustly to changes in ambient CO_2_ levels, independent of the concentration. Adding either air or CO_2_ to the arena triggered short-term startle responses, presumably because of a change in airflow. After the startle, CO_2_ exposure generally altered tick behavior for the remainder of the experiment. By contrast, during air puffs, the ticks typically resumed the behavior they had shown before the puff.

### CO_2_ elicited behavioral reactions are state-dependent

We noted that a significant proportion of the animals did not walk and were stationary already in the control condition. It is well-known that an animal’s behavioral state can influence the assessment of a sensory stimulus and associated behavioral responses. For example, fruit flies (*Drosophila melanogaster)* are attracted to CO_2_ while flying but repelled by it when walking (Wasserman et al., 2013). We thus hypothesized that a similar state dependency might exist in ticks and that the way ticks respond to CO_2_ depends on whether the animals are walking or not.

We used ticks that molted 10 to 18 weeks before testing, a time that *Ixodes scapularis* easily survives. This time is also needed to regain interest in potential hosts, as this is greatly reduced for a few weeks after molting (Gray et al., 2016). Ticks can survive long periods without a blood meal. Yet, energy resources are limited and ticks avoid unnecessary activities and intermittently rest or become dormant (Gray et al., 2016). However, non-walking animals were not dormant or unresponsive in our experiments. Resting ticks frequently waved their forelegs at irregular intervals, indicating they were still sampling their environment. We refer to these quiescent but responsive animals as stationary.

To test whether the animal’s physiological state affected their response to CO_2_, data from walking and stationary ticks were separated based on the behavior displayed in ambient air (Figure 2). We analyzed changes in walking speed and foreleg movement two minutes before and after the start of treatment. We report percentage values rather than animal numbers because we noted a steady decrease in walking animals during the three-hour experiments while the number of stationary animals increased (Figure 2A, *middle column: 80% walking at the beginning, 42% at the end*). Since treatment order was not randomized in these experiments, the shift from walking to stationary is most likely caused by incipient dehydration over experiment time. However, CO_2_ responsiveness was not affected by this shift.

**Figure 2:**
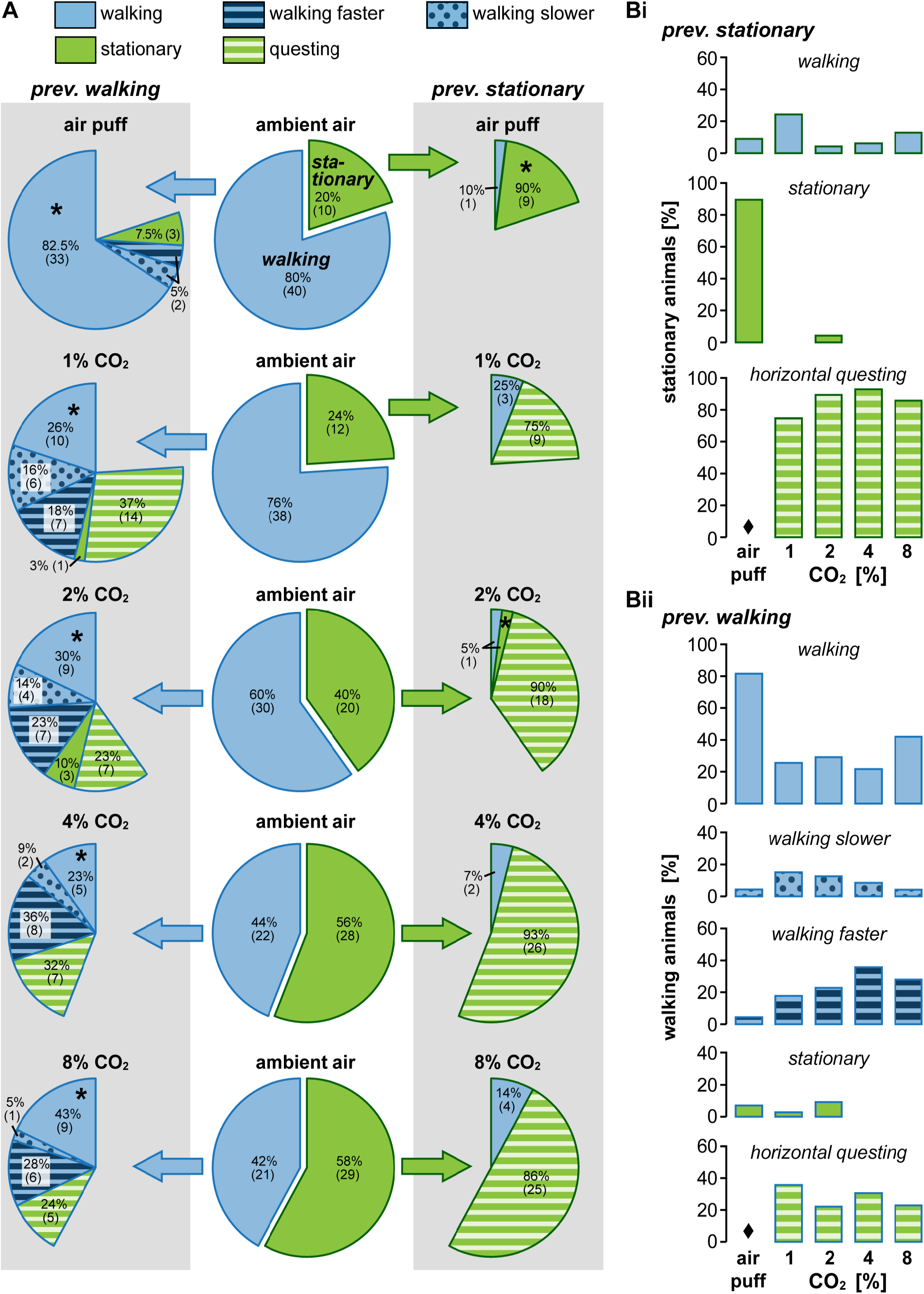
Changes in behavior are state-dependent - walking animals tend to increase their walking speed while stationary animals start to wave their forelegs. **(A)** Manual video analysis of behavioral changes before and during treatment for the same 50 animals as in Figure 1, sorted by treatment. Middle column (white background): Pie charts showing the percentage of walking (blue) and stationary animals (green) before treatment in ambient air. Left and right columns (gray background): behaviors displayed by previously walking (left) and stationary animals (right) during the treatment (air puff or CO_2_). Behaviors are color-coded, with different walking behaviors shown in blue and stationary behaviors in green. The numbers denote the percentage of animals with N numbers in parenthesis exhibiting a particular behavior. The sum of the wedges on either side is 100%, regardless of whether it is a full circle. Asterisks highlight animals that showed no behavioral change in response to the treatment (walking → walking; stationary → stationary). **(B)** Same data as shown in A but here sorted by behavior for (i) previously stationary animals and (ii) previously walking animals. ♦ highlights that none of the animals tested (walking and stationary) started to quest in response to the air puff.

Figures 2A and 2B show the same data, visualized differently (A: sorted by treatment; B: sorted by behavior). Exposure to CO_2_ clearly induced behavioral changes in stationary and walking animals. Surprisingly, CO_2_ did not induce robust walking in stationary animals. Most stationary animals remained stationary when exposed to CO_2_ but started to wave their forelegs (Figure 2A, green-striped wedges in right column). Host-seeking *Ixodes scapularis* often quest by climbing a patch of grass, stretching out, and waving their forelegs while waiting for passing hosts (Keirans et al., 1996). Because of the similarity to the natural behavior, we refer to increased foreleg waving and posture changes as questing. Each CO_2_ concentration tested elicited questing in stationary animals, with a peak at 4% CO_2_ (93%) and a minimum at 1% CO_2_ (75%). Ticks often remained in the questing position until CO_2_ levels were reset to ambient air levels at the end of an experimental block. This long-lasting foreleg waving was exclusively observed during CO_2_ exposure and differed from the brief startle responses. None of the animals started to quest in response to the air puff (visualized with ♦ in Figure 2Bi). Only a few stationary animals began to walk (Figure 2A, blue wedges in right column), with the highest percentage (25%) at 1% CO_2_ (Figure 2Bi).

The response of walking animals was markedly different: Most walking animals continued to walk in the presence of CO_2_ (Figure 2A, sum of blue colored wedges in left column), walking speeds differed substantially between the air puff and CO_2_ exposure. During the air puff, most ticks showed no change in walking speed, but in CO_2_, most ticks either slowed down or sped up. The highest percentage of slower walking animals (blue dotted wedges) was observed at 1% CO_2_ (16%, Figure 2Bii). Faster walking (blue-striped wedges) was most pronounced at 4% CO_2_ (36%), suggesting that higher CO_2_ concentrations promote faster walking. Independent of the CO_2_ concentration, approximately one-third of the walking animals became stationary, and most of these stationary animals started to quest (green-striped wedges in the left column). This starkly contrasted the responses to air puffs, where only a tiny percentage of the animals stopped walking, and those did not quest (* in Figure 2Bii). Given that walking speed changed when CO_2_ was applied, we suspected that walking direction and the number of directional changes were also affected. However, we found no consistent pattern in number of directional changes, turning direction and magnitude, angular acceleration, and walking direction, neither in the control conditions nor in CO_2_ (data not shown). In summary, CO_2_ behavioral reactions were state-dependent in that stationary animals started to quest, while walking animals continued to walk but at different speeds. The number of animals that increased their walking speed rose with the CO_2_ concentration and reached its maximum at 4% CO_2_. Questing in stationary animals was pronounced at all tested CO_2_ concentrations but strongest at 4%.

### CO_2_ is sufficient to elicit foreleg waving

Although not well documented in the scientific literature, certain ticks stretch out and wave their forelegs when breathed on. However, breath contains many potent stimulants besides CO_2_, so it is unclear whether CO_2_, another respiratory compound, or the blend itself induces foreleg waving. McMahon and colleagues (2002) found that *Amblyomma variegatum* responds strongly to exhaled air, but CO_2_ alone failed to mimic this response. While for *Amblyomma variegatum*, the breath blend may be more influential than CO_2_ alone, our data show that CO_2_ is sufficient to trigger foreleg waving in *Ixodes scapularis*. To characterize CO_2_-induced changes in foreleg waving, we performed single animal experiments by exposing 8 *Ixodes scapularis* individually to 4% CO_2_. Ticks were tethered to a metal rod and positioned on an air-supported ball (Figure 3A). The ball acted as an omnidirectional treadmill, allowing the animals to walk without changing position. Like before, an air puff was used as a negative control, and animals were video-monitored from above.

**Figure 3:**
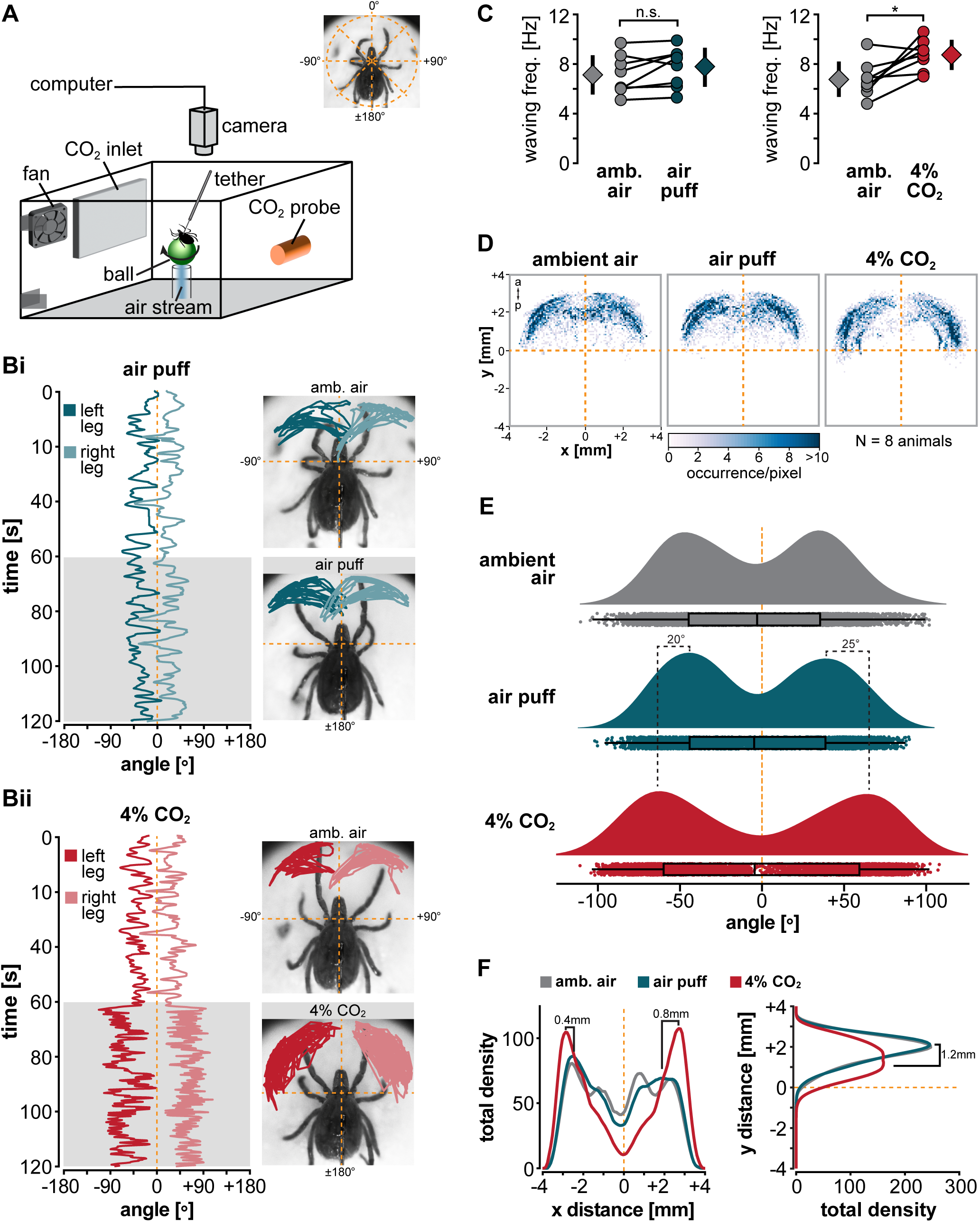
CO_2_ elicits foreleg waving. **(A)** Schematic of the single-animal tick-on-a-stick assay. Rigid-tethered animals were video monitored while walking on a floating ball. Inset on the top illustrates the polar coordinate system used for data analysis. **(B)** Space-time diagrams depicting the angular positions of the left (negative values) and right (positive values) foreleg tip as a function of time. Example traces for one adult walking female. Measurements in (i) and (ii) are from the same animal. Leg movement was monitored over 60 seconds in ambient air (white background) before the treatment started (gray rectangle, 60 seconds; (i) air puff, (ii) exposure to 4% CO_2_). Sampling rate was 5 frames/s. The dashed yellow lines depict the midline of the animal. The images on the right show the x-y positions of the left and right foreleg tips in the polar coordinate system (top: ambient air, below: treatment; 60 s, respectively). Supplementary Figure S3 shows the data for each individual animal tested. **(C)** Comparisons of waving frequency for all 8 animals tested (4 males and 4 females). Dots represent the average waving frequency for individual animals, measured over 60 seconds. Diamonds depict the population means ± SD as vertical bars. *Statistics: paired t-test; n*.*s. = no significant difference, t(7) = -1*.*75, p = 0*.*12; * significant different, t(7) = -3*.*23, p = 0*.*014*. **(D)** Colormaps showing the positions of the left and right foreleg tips as a function of the distance from the animals’ midline (dashed yellow lines) for the 8 animals tested. Shades of blue represent the number of occurrences per pixel (100 x 100 pixels, 0.08 mm/pixel). Body orientation is marked with a (anterior) and p (posterior). **(E)** Raincloud plots showing the angular distribution for the left (negative angles) and right (positive angles) foreleg tip for all 8 animals tested. Each chart shows the probability density estimate (colored areas), the corresponding box plots (median with 25/75% quartiles), and raw observations (dots) for 4800 foreleg positions (120 s recordings at 5 frames/s for 8 animals). The Y-axis is scaled the same for each chart and shows the density distribution of the data points in arbitrary units. The area of each density distribution is equal. Changes in height correspond to changes in density. The more frequently an angle occurs, the higher the density curve. **(F)** Distribution of the density of foreleg tip positions as a function of the distance from the animals’ midline (dashed yellow lines) for the x-axis (left) and y-axis (right). The mean values of foreleg tip positions are shown for all 8 animals tested for 100 x 1 pixels (x and y axis, respectively).

We analyzed the angular position of the forelegs over 120 seconds (60 seconds in ambient air and CO_2_, respectively) by placing a virtual coordinate system through the basal part of the animal’s mouthparts (Figure 3A, inset). Figure 3B shows an example of this analysis for one adult walking female during an air puff (i) and exposure to 4% CO_2_ (ii). The plots on the left show the angles of the forelegs over time, while the images on the right show the position of the forelegs in relation to the animal. Supplementary Figure S3 shows the forelimb movement of all 8 animals tested. During CO_2_ exposure, two noticeable changes occurred: the waving frequency increased, and the animal spread its forelegs further. This is evident by larger oscillations and angles in the red traces in Figure 2Bii.

To quantify the effect of CO_2_ on foreleg waving, we measured waving frequency over 60 seconds before and after the start of treatment (Figure 3C). Waving frequency during the air puff did not differ significantly from ambient air (*ambient air* (mean ± SD): 7.13 ± 1.59 Hz; *air puff*: 7.74 ± 1.58 Hz; p>0.05). By contrast, during 4% CO_2_ exposure foreleg waving frequency increased in 7 of the 8 animals tested, and the increase was significant compared to ambient air (*ambient air*: 6.78 ± 1.42 Hz; *4% CO*_*2*_: 8.74 ± 1.22 Hz; p<0.05). The spread of the forelegs was also consistent. The heatmaps in Figure 3D depict the position of the forelegs for all 8 animals tested. In ambient air and during the air puff, the animals moved their forelegs within a range of approximately ±90° from the midline, with the highest density at ±40° to ±50° (Figure 3E). In the presence of 4% CO_2_, the range of motion increased to approximately ±100° from the midline, with the highest density at about ±70°. On average, foreleg motion shifted 0.4 to 0.8 mm outwards from the midline and 1.2 mm posteriorly (Figure 3F).

We noted that none of the tested animals were stationary in these experiments. All animals walked continuously or intermittently in control ambient air and during treatments. The air-supported ball appeared to have motivated the animals to walk, which was beneficial to our study because we could determine that CO_2_ induces foreleg waving in walking ticks, too. We also observed that animals did not consistently use their forelegs for locomotion but instead appeared to use them for active sensing. This is based on the observation that the gait often switched from eight-legged walking in ambient air to six-legged walking with extended and rapidly moving forelegs in CO_2_. Active sensing is a common phenomenon in arthropods akin to sniffing in mammals. In insects, movement of the usually bilaterally symmetric sensory appendages during active sensing is often coordinated and coupled to leg movements (Dürr et al., 2001; Horseman et al., 1997). We noticed, however, that movements of the left and right forelegs were not strongly synchronized in our experiments. There were brief synchronized phases in which the left and right legs moved approximately parallel towards each other (left-right waving) or in opposite directions (inward-outward waving) and uncoordinated phases in which the left and right legs moved independently (Supp. Figure S3). This was the case in all animals tested in ambient air, during the air puff, and CO_2_. This suggested that, at least at times, the coupling between the left and right foreleg was weak or absent and that legs can be independently controlled for active sensing. However, our assay could not detect 3-dimensional movements of the legs. Whether or not the movement patterns of *Ixodes’* forelegs are goal-directed and crucial for mapping the olfactory space remains to be determined.

### The Haller’s organ is not necessary for CO_2_ detection

The mechanisms of tick chemoreception are unknown, but they are mainly ascribed to the foreleg HO (Carr et al., 2017; Gebremedhin et al., 2023). The HO is unique to ticks and mites and presumed to function like the insect antennae. The structure and organization of the HO varies across tick species (Josek et al., 2018). Generally, the HO is located on the dorsal surface of the tarsus of each foreleg. It includes four functionally and structurally different parts: a capsule, an anterior pit, the distal knoll, and a post-capsular area. Each part contains a definite number of structurally different sensilla. The morphology of the HO in adult *Ixodes scapularis* is insufficiently understood. The only available examples come from transmission electron microscopy of the HO of *Ixodes persulcatus* (Leonovich, 1977) and light microscopy studies of *Ixodes ricinus* (Leonovich and Belozerov, 1992). In these species, the capsule contains 7 wall-pored sensilla, which are presumed olfactory based on their morphology. The anterior pit contains 6 sensilla. One is wall-pored and innervated by 6 bipolar sensory neurons. It is unclear how many sensilla the posterior capsule of the genus *Ixodes* contains, as it is covered by cuticle except for a slit and is difficult to access. However, the anterior pit of *Amblyomma variegatum* bears 7 sensilla. Three of these sensilla are wall-pored and CO_2_-sensitive (Steullet and Guerin, 1992).

Although not previously described in detail, we are not the first to notice that CO_2_ elicits foreleg waiving in certain tick species. It is, therefore, widely believed that the HO is the main sense organ for detecting CO_2_. To our knowledge, however, no study has carefully tested this hypothesis. We thus tested ticks with disabled HO for CO_2_ responses by covering the foreleg tarsi with wax. From an experimental standpoint, sufficient covering requires a substance that changes from liquid to solid in seconds, can be applied in multiple layers, has a sufficiently low viscosity without dripping, and is reasonably flexible after hardening. Care also needs to be taken to not cause damage to muscle or nerve tissue through excessive heat or harsh chemicals. Resins such as UV-curable adhesives and epoxies cure quickly but do not adhere well to the tick’s cuticles for an extended period. Fast-curing dental silicones work well, but the curing time is too slow, at 2-3 minutes per layer, since multiple layers are required to completely cover the long wall-pored sensilla of the HO capsule. We ended up using low-melting point wax since it fulfilled all requirements.

Figure 4A shows an adult female with covered HO. First, we confirmed that the wax was impermeable to CO_2_. For this, a CO_2_ probe was covered with a thin layer of wax (Supp. Figure S4D), about the same thickness as the wax applied on the forelegs. The covered probe was placed inside the behavior arena, and CO_2_ was added for 2.7 and 6 seconds. If the wax is impermeable to CO_2_, measurements should stay the same once CO_2_ is present. Figure 4C shows that this was the case. CO_2_ concentration in the arena increased only by 0.015% (150 ppm) on average for 2.7 and 6 seconds of CO_2_ application, respectively. This amount is within the measurement uncertainty of the probe. For comparison, 2.7 s CO_2_ application resulted in a concentration of 4% and 6 s in 10% when the probe was not covered with wax.

**Figure 4:**
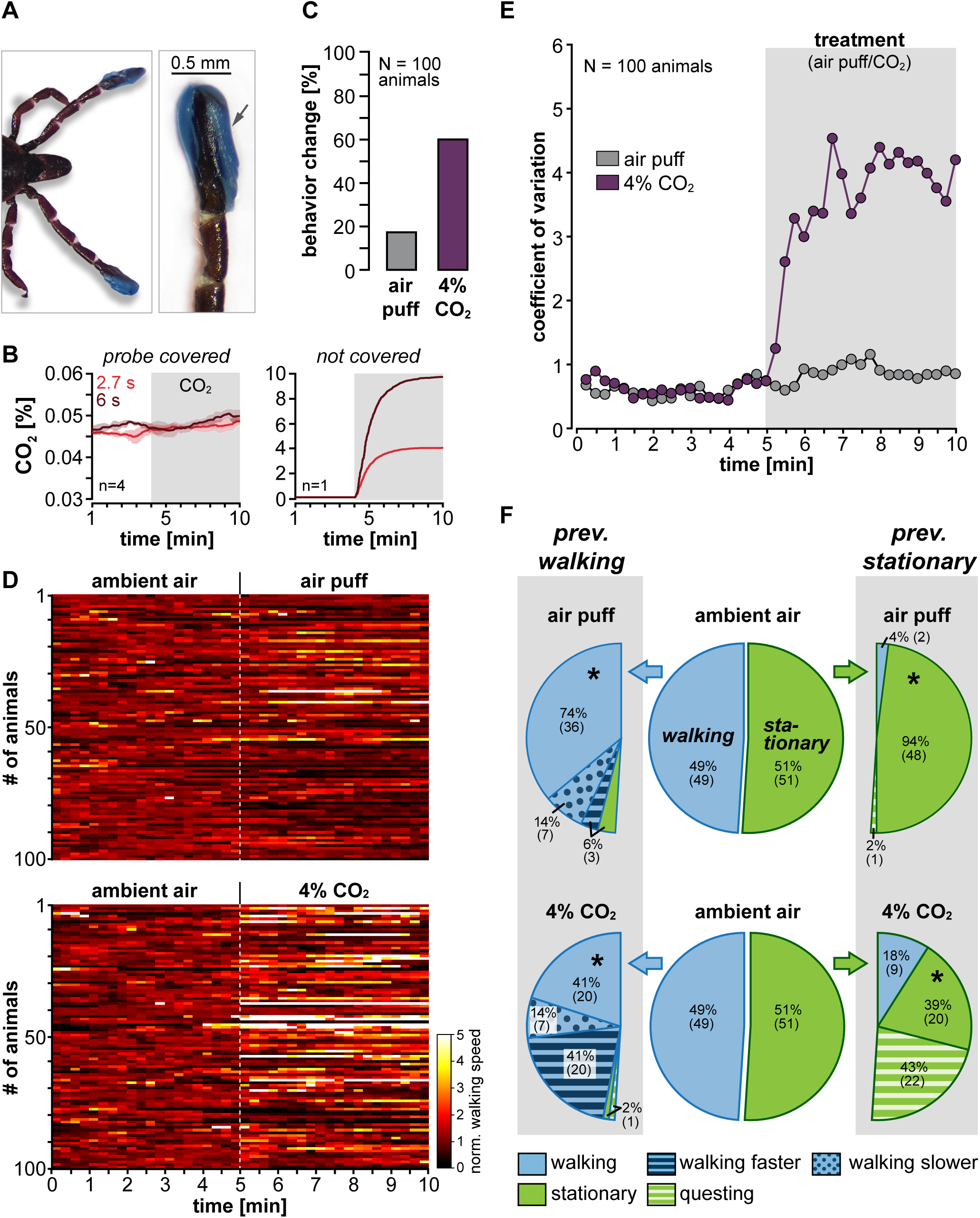
Disabling the Haller organ does not eliminate CO_2_ responses. **(A)** Photos showing the covering of the foreleg tarsi and the Haller’s organ with blue-colored low melting point wax. Care was taken to completely cover all sensilla, including the long distal wall-pored sensilla on the Haller’s organ capsule (arrow in the right image). **(B)** Left: Comparison of wax-covered CO_2_ probe measurements for two different CO_2_ inlet times (6 and 18 s). Mean values from 4 measurements ± SD (shaded area) are shown. Right: Comparison of non-wax covered CO_2_ probe measurements for 2.7 and 6 s CO_2_ inlet (single trial). The gray rectangle illustrates the presence of CO_2_. **(C)** Analysis of behavioral changes of 100 adult *I. scapularis* (50 males and 50 females) with covered Haller’s organ during exposure to an air puff and 4% CO_2_. Data were obtained by manual video analysis and include changes in walking speed and foreleg/body position. See Figure 1B for a comparison to animals with intact Haller’s organ. **(D)** Analysis of normalized walking speed before (ambient air) and during exposure to an air puff (top) or 4% CO_2_ (bottom). Same 100 animals as in panel B. For a more detailed description, see Figure 1D legend. **(E)** Coefficient of variation for the data presented in panel C. Dots represent mean values. The gray rectangle illustrates the start and duration of the treatment (air puff or 4% CO_2_). **(F)** Pie charts showing the percentage of exhibited behaviors for 100 animals with covered forelegs before (ambient air) and during exposure to 4% CO_2_. Animals were sorted based on their dominant behavior in ambient air (walking or stationary). For more details, please refer to Figure 2A legend.

We then tested *Ixodes*’ responses in the multi-animal walking arena (Figure 1A) to CO_2_ application with the HO covered with wax. One hundred ticks in groups of 5 were tested. We only tested responses to 4% CO_2_, since *Ixodes scapularis* did not show a strong concentration preference (Figure 1C). Air puffs were used as a negative control, and the order of CO_2_ and air puff was randomized. We first noted that ticks with covered HO rarely startled at the beginning of the treatment, evident by the missing dark band at the treatment onset (Supp. Figure S4A). Only 8% of the 100 ticks with covered HO decreased their walking speed at the onset of the air puff, and 6% for the 4% CO_2_ treatment (Supp. Figure S4C). For comparison, the air puff triggered startle responses in 30% of the animals with intact HO and in 24% for the 4% CO_2_ treatment. The decrease in startle responses shows that the wax coating was successful, and that the HO appears to detect changes in airflow that prompt startle responses.

Like for ticks with intact HO, the air puff did not strongly alter *Ixodes’* behavior throughout the trial. Only 18% of the animals with covered forelegs showed changes in behavior (Figure 4C). Surprisingly, however, 60% of the animals responded to 4% CO_2_, although the HO was covered. Likewise, a comparison of normalized walking speeds in ambient air and during treatment revealed that walking speeds increased in response to 4% CO_2_. The increase in walking speed is visible in the shift to lighter colors in the heatmaps (Figure 4D) and the increase in CV during the 4% CO_2_ treatment (Figure 4E). Additionally, we noted that ticks with covered HO walked significantly slower than with intact HO, even in ambient air (*intact HO*: 1.0 ± 0.7 mm/s, N=22; *covered*: 0.5 ± 0.4 mm/s, N=49; t-test: t(69) = 4.712, p < 0.001). However, the wax application did not interfere with Ixodes’ ability to walk. Walking was just slower. Although walking speed in ambient air was initially decreased in ticks with covered HO, this difference disappeared during CO_2_ exposure. Most ticks that changed their walking speed in the presence of 4% CO_2_ started to walk faster (Figure 4F, blue-striped wedges, left panel). They reached speeds similar to ticks with intact HO (*intact HO*: 1.6 ± 1 mm/s, N=13; *covered*: 1.5 ± 0.7 mm/s, N=40; t-test: t(51) = 0.247, p = 0.806).

Covering the HO had further effects: First, fewer animals started to quest when the HO was covered. Only two walking ticks became stationary, with one starting to quest during 4% CO_2_ exposure (Figure 4F, green colored wedges, left panel). Questing was still present in stationary animals, albeit less prevalent than in animals with intact HO (green-striped wedges, right panel). Concurrently, the percentage of stationary animals that remained stationary without questing (green wedges, right panel) was increased. Second, questing duration was reduced. Untreated animals often quested until the CO_2_ concentration was restored to ambient air levels at the end of an experimental block (5 minutes and longer). In contrast, animals with covered HO only quested for 157 ± 67 seconds on average (N=23 ticks). Third, the latency between the start of the 4% CO_2_ treatment and the onset of behavioral reaction was significantly increased. Untreated ticks with intact HO showed behavioral changes after 3.8 ± 2.5 sec (N = 45 ticks). In comparison, ticks with covered HO showed behavioral changes after 17.1 ± 7.2 s (N = 60 ticks, *t test: t(103) = -11*.*899, p < 0*.*001 compared to 4% CO*_*2*_ *intact HO*).

We excluded that the additional weight of the wax caused the decrease in questing by applying a comparable amount of wax to the patella proximal on the forelegs (Suppl. Figure S5). The additional weight on the patella did not strongly impact CO_2_ reactions. A similar number of ticks with wax on the patella responded to 4% CO_2_ as untreated ticks with intact HO (84% versus 90%). In addition, all stationary animals that remained stationary started to quest and the onset of behavioral changes was not statistically different from untreated animals *(wax on patella*: 4.1 ± 3.2 s, N = 42 ticks, t test: t(85) = -0.436, p = 0.664 compared to 4% CO_2_ intact HO).

We conclude that the HO is not necessary for CO_2_ detection. However, CO_2_ induces foreleg waving, supporting that CO_2_ acts as an activator and primes the animals to quest for additional sensory cues. Fewer ticks started questing, and questing time was reduced when the HO was covered. This suggests that the animals lose interest in CO_2_ more quickly when no active sensory input from the HO exists.

### Animals with amputated forelegs still respond to CO_2_

Our data contradict the assumption that *Ixodes scapularis* HO is the primary CO_2_ detector. Therefore, we performed an additional control experiment, amputated the foreleg tarsi and the HO for 50 ticks, and tested responses to 4 % CO_2_ and the air puff (Figure 5A). The amputation affected *Ixodes scapularis* behavior, especially in the smaller males. Seven of the 25 amputated males died within 24 hours after amputation, while all females survived. As before, ticks with amputated forelegs walked slower in ambient air (0.4 ± 0.5 mm/s, N=22 ticks) than walking animals with intact HO. Regardless of the animals’ potentially impaired condition, 58% of the animals responded to 4% CO_2_ either by changing their walking speed or starting to quest (Figure 5B-D). However, walking animals that continued walking in CO_2_ did not reach the same average speed as ticks with covered HO or untreated animals (0.9 ± 0.3 mm/s in comparison to ∼1.5 mm/s, N=14). Overall, more animals decreased their walking speed in response to CO_2_ (Figure 5E, blue-dotted wedges, left panel), most likely as a result of the amputation. Similar to ticks with covered HO, fewer animals with amputated forelegs started to quest in response to 4% CO_2_ (Figure 5E, green-striped wedges), and questing time was reduced (169 ± 46 s, N=13 ticks). In addition, the latency between the start of the 4% CO_2_ treatment and the onset of behavioral reaction was significantly increased compared to 4% CO_2_ exposure of ticks with intact HO (17.9 ± 7.2 s, N = 29 ticks, *t test: t(72) = -12*.*065, p < 0*.*001 compared to 4% CO*_*2*_ *intact HO*).

**Figure 5:**
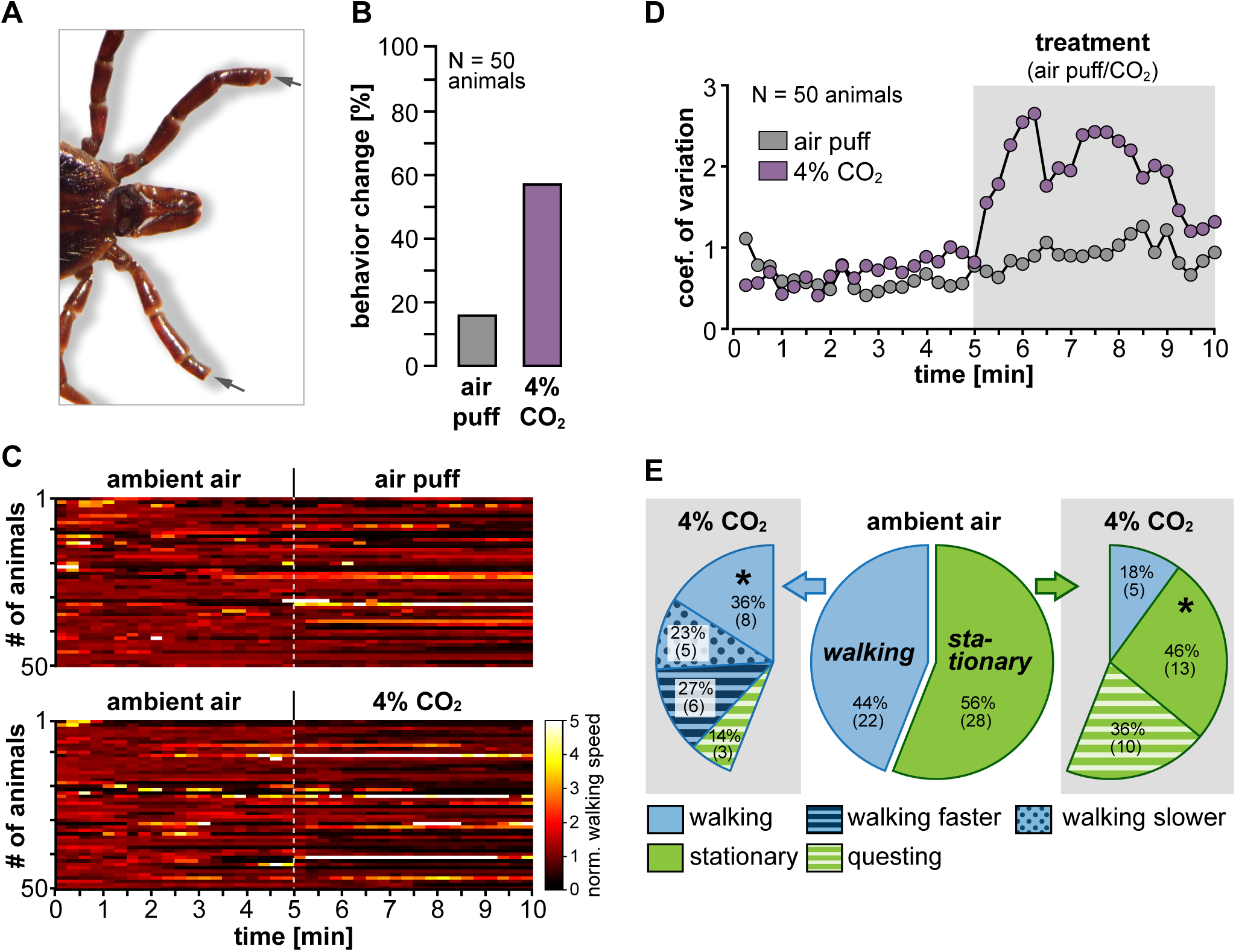
Ticks with amputated Haller’s organ still respond to CO_2_. **(A)** Photo showing the cutting site on the foreleg tarsi (arrows). **(B)** Manual video analysis of behavioral changes of 50 adult *I. scapularis* (25 males and 25 females) with cut Haller’s organ during exposure to an air puff (=negative control) or 4% CO_2_. **(C)** Analysis of normalized walking speed before (ambient air) and during exposure to an air puff (top) or 4% CO_2_ (bottom). Same 50 animals as in panel B. For a detailed description, see Figure 1D legend. **(D)** Coefficient of variation for the data presented in panel C. Dots represent mean values. The gray rectangle illustrates the start and duration of the treatment (air puff or 4% CO_2_). **(E)** Pie charts showing the percentage of exhibited behaviors for the same 50 animals as in panel B with cut foreleg tarsi before (ambient air) and during exposure to 4% CO_2_. For more details, see Figure 2A legend.

In search of the structure mediating CO_2_ detection, we covered the palpal organ with wax in a subset of experiments in addition to the HO. The palpal organ is a sensory structure on the tick mouthparts, presumed to be involved in chemosensation. Several multi-innervated sensilla have been identified in *Amblyomma americanum* that either have an elaborate pore canal system or a single slit opening at the tip (Foelix and Axtell, 1971). However, covering the palpal organ and its sensilla did not prevent *Ixodes* from reacting to CO_2_. 65% of the 20 animals tested changed their walking speed or started to wave their forelegs in response to 4% CO_2_ (Supp. Figure S6). This suggests that other sensory structures, besides the palpal and HO mediate CO_2_ responses in *Ixodes scapularis* that still need to be identified.

Taken together, ticks with amputated and covered HO reacted similarly robust to CO_2_, providing further evidence that the HO is not necessary for CO_2_ detection.

## DISCUSSION

In summary, adult male and female *Ixodes scapularis* respond robustly to CO_2_ in the tested concentration range of 1 to 8%. Notably, CO_2_ responses were maintained even when the HO was disabled, either through covering it with wax or by amputating the foreleg tarsi.

### CO_**2**_ can serve as a long-distance host signal

We did not find a clear concentration preference. While the highest number of behavioral responses was observed during exposure to a breath-like concentration of 4% CO_2_, all CO_2_ concentrations elicited robust behavioral responses with only marginal differences (Figure 1C).

The lack of a strong CO_2_ concentration preference may seem counterintuitive. However, a concentration preference is not required to locate a CO_2_ source. Odors are not distributed in a laminar manner in the air but rather as turbulent plumes of fine odor filaments with defined edges (Rigolli et al., 2022). The spatiotemporal pattern of these filaments depends on the distance from the odor source. Concentrations are typically highest at the origin and decreases with distance, while the odor plume becomes increasingly patchy and intermittent. While turbulent wind currents in the environment carry odorants from a source to an animal’s olfactory organ (e.g., antenna, nose, forelegs), small-scale currents near the organ surface and molecular diffusion carry odorants to the olfactory receptors (Ache et al., 2016). Many animals actively interact with these small-scale currents, for example by sniffing or increased movement of sensory structures such as antennae or forelegs, also known as active sensing (Crimaldi et al., 2022).

Given the turbulent structure of odor plumes, detecting and staying within the edges of an odor plume is a more reliable detection strategy than using the concentration gradient itself. To detect edges, noticing CO_2_ already at low concentrations is essential. Mosquitoes, for example, track their hosts over long distances - up to 50 meters in some cases – by following host-emitted signals such as CO_2_ (Dekker et al., 2005; Healy and Copland, 1995). Many mosquito species can detect even small increases in the ambient CO_2_ concentration. In *Aedes aegypti* mosquitoes, CO_2_-sensitive maxillary palp sensilla neurons increase their action potential firing rate as soon as the CO_2_ concentration is slightly above ambient levels (Grant and O’Connell, 2007). Whether *Ixodes scapularis* have a similarly low CO_2_ sensitivity is unknown. However, a study using the tropical bont tick *Amblyomma variegatum* has demonstrated that sensilla in the HO show threshold CO_2_ responses between 0.002% and 0.1% (Steullet and Guerin, 1992). This suggests that, at least in principle, this tick species can detect the CO_2_ plume of relatively distant hosts. Whether this is also the case for *Ixodes scapularis* remains to be tested. However, in our experiments, already 80% of the animals tested reacted to 1% CO_2_ exposure (Figure 1C), suggesting that detection of CO_2_ at concentrations below 1% is likely. Our results also indicate that CO_2_ can serve as a long-distance host signal and that hungry adult *Ixodes scapularis* become aware of approaching hosts early on. Detecting hosts from a greater distance is beneficial as it gives the animals more time to gather additional information and decide if attaching is rewarding.

### CO_**2**_ activates host-seeking behavior in *Ixodes scapularis*

Host-seeking is not a single behavior but rather a sequence of behaviors. A typical host-seek ing sequence in endo- and ectoparasites alike is 1) activation of the animal, 2) orientation towards the potential host, and finally, 3) host selection and invasion. The expression of each of these phases varies between different animal species. Thus, it is sensible to assume that orientation behaviors differ between ticks that primarily quest for hosts and ticks that actively hunt. Our data show that CO_2_ stimuli activate *Ixodes scapularis*, and initiates behaviors that resemble host-seeking. Specifically, we identified two behavioral responses that depended on the state of the animal and can readily be assessed in laboratory conditions: walking ticks primarily increased their speed, mirroring a faster approach to a host during active hunting. In contrast, stationary ticks raised their forelegs and began to wave (Figure 2). Foreleg waving strongly resembles *Ixodes scapularis’* questing behavior in their natural habitat. It also mirrors the antennae and forelimb movements of other arthropods during active sensing. Active sensing is widespread in animals and is used, among other things, to recognize additional sensory cues and to sample the environment.

CO_2_, by itself, is an ambiguous sensory stimulus. It is a by-product of plant respiration, decomposition of plant and animal matter, and the burning of oil, coal, and gas. Thus, a CO_2_ plume encountered in the environment can have vastly different behavioral relevance. Approaching a non-host CO_2_ source can endanger the animal or, at best, waste already scarce energy resources. Information about the context of the CO_2_ presence is thus essential to make optimal decisions. Breathing animals provide visual targets, create thermal gradients, and release hundreds of compounds from the skin and in the breath, all contributing to a species-specific multi-modal sensory stimulus. It is thus likely that ticks use strategies similar to parasitic nematodes and mosquitoes (Hallem et al., 2011; McMeniman et al., 2014; Vinauger et al., 2019), where CO_2_ enhances the attraction to other sensory stimuli, such as host-specific odors and body temperatures resembling vertebrate body heat.

### CO_**2**_ may promote active sensing to detect tactile stimuli

The forelegs of ticks carry the HO, a multisensory organ implicated in olfaction (Carr et al., 2017; Josek et al., 2021) and thermotaxis (Carr and Salgado, 2019; Mitchell et al., 2017) in certain tick species. Although direct evidence in *Ixodes scapularis* is lacking, it is conceivable that there are also mechanosensory sensilla on the foreleg tarsi. We suggest that the CO_2_-evoked waving of the forelegs in stationary and walking animals (Figures 2 & 3) is a form of active sensing and allows the detection of additional (multimodal) stimuli. In fact, we observed that walking animals that already moved their forelegs during walking significantly increased the speed and range of waving movements in CO_2_. This is consistent with the notion that the increased waving of the forelegs is based on the need to increase the capture rate of odorants, for example. We also observed that ticks occasionally erected their entire body while waving their forelegs. To effectively detect potential air-born or thermal stimuli, it should not make a difference for the animals whether they are erected or not. Most odor-tracking arthropods do not erect their bodies as it is an energetically costly behavior and may expose the animal to predation. However, erecting the body is beneficial for getting closer to a potential host and obtaining mechanosensory information when the host is within tangible reach. We thus postulate that after CO_2_ exposure, ticks are more motivated to seek mechanoreceptive touch stimuli that enable them to attach to hosts. This idea is consistent with our observation that after exposure to CO_2_ ticks were more likely to grab onto a paintbrush that was brought into their immediate proximity. On the other hand, animals not exposed to CO_2_ crouched or did not react. While we did not quantify these observations, they suggest that CO_2_ activates and promotes mechanosensory host-seeking in *Ixodes scapularis*.

Mechanosensation in ticks is vastly unexplored, and to our knowledge, there are no examples from other animal species that CO_2_ primes tactile stimulation. However, ticks belong to the class *Arachnida* along with spiders, scorpions, mites, and harvestmen, many of which have highly developed mechanosensory systems for detecting prey and predators, as well as for proprioception and communication with potential mates (Barth, 2002). The exoskeleton of many arachnids is covered with mechanosensory touch hairs or sensilla. The cuticula and legs of *Ixodes scapularis* are also covered with hair-like sensilla. It is reasonable to assume that many of these sensilla are mechanically sensitive and involved in controlling body and joint positions. However, we suspect that mechanosensory sensilla on the forelegs are also essential for host attachment. An electron microscopic examination of the foreleg tarsi of the cattle tick *Rhipicephalus microplus* (formerly *Boophilus microplus*) has shown that the morphology of sensilla surrounding the HO resembles that of mechanosensors (Waladde, 1977), supporting our hypothesis.

### *Ixodes scapularis* has two separate CO_**2**_-detection pathways

So far, the only CO_2_-sensitive structures identified in ticks have been found in the HO capsule of the tropical bont tick *Amblyomma variegatum* (Steullet and Guerin, 1992). Two of the seven capsule sensilla in *Amblyomma variegatum* are sensitive to changes in CO_2_ concentration. CO_2_ elicits opposite responses in the afferent neurons of the two sensilla. In one sensillum, neurons decrease their action potential firing rate as soon as CO_2_ levels are about 0.002% above ambient, while neurons in the other sensillum are less sensitive and increase their firing rate at CO_2_ levels above 0.1%. Apart from this study, no other examples of CO_2_-sensitive sensilla in other tick species are available and it is unclear if the HO of *Ixodes scapularis* bears similarly CO_2_-sensitive sensilla.

Our results convincingly demonstrate that *Ixodes scapularis* still responds to CO_2_ even when the HO is disabled (Figures 4 & 5). In fact, the CO_2_ responses were only slightly affected by disabling the HO. Slightly fewer animals responded and the response onset was delayed in comparison to ticks with intact HO. This result was surprising and caused us to carry out extensive control experiments to rule out experimental artifacts. We doubled the number of animals tested and determined that the wax used to cover the forelegs is CO_2_ impermeable (Figure 4B). We also ruled out that the wax application itself might have affected the animals’ behavior (Supp. Figure S5). We even amputated the forelegs and the HO and the animals still showed robust responses to CO_2_. Taken together, our data thus demonstrate that the HO is not necessary for CO_2_ detection and the subsequent initiation of host-seeking behavior.

However, this does not imply that the HO is not involved in CO_2_ detection. Our data indicate that the HO might be able to detect CO_2_. When we examined the startle responses that occurred immediately after treatment onset, we found that they were not independent of the CO_2_ concentration. Although ticks showed startle responses during both air and CO_2_ stimuli, the number of responding animals was highest at the lowest CO_2_ concentration (1%, Supp. Figure S2B). This suggests that the presence of air movement is recognized, but that low levels of CO_2_ enhance the startle response. Furthermore, our data show that startle responses are mediated by HO. Startle responses were almost absent when the HO was disabled (Supp. Figure S4C). Consequently, CO_2_ sensitive structures in the HO appear to contribute to the startle responses and mediate a rapid detection of CO_2_ stimuli - particularly at lower CO_2_ concentrations. This hypothesis is consistent with the observation that the latency between CO_2_ onset and behavioral responses in ticks with disabled HO was significantly delayed than in animals with intact HO. Furthermore, the startle responses support that one of the functions of the HO is to actively sense CO_2_ plumes and their edges and implies that the HO of *Ixodes scapularis* contains CO_2_-sensitive afferent neurons that respond to low concentrations.

Our main finding is that there must be a separate CO_2_-detection pathway that initiates the orientation phase of the host-seeking behavior, i.e., the questing and faster walking. While we could not identify the CO_2_ detectors that initiate the behavioral responses, we can rule out the Haller’s and palpal organs (Figures 4 & 5). A possible candidate is the paired spiracle plate located on the ventrolateral surface of the tick body directly posterior of each hindleg coxa. The spiracle plates are the main respiratory organ in ticks and important for gas exchange (Soneshine and Roe, 2014). Regardless of the structure that mediates the CO_2_ detection, it is most likely conveyed via passive sensing. In comparison to active sensing where energy is invested to sample the environment, passive sensing is cost-effective. However, the disadvantage of passive sensing is that it depends on the availability and quality of the stimulus. It typically has lower resolution and accuracy, and is often inadequate for determining target distance, speed, and direction. This fits our observation that the behavioral responses were independent of CO_2_ concentration, and it is consistent with the idea that there are two parallel CO_2_ detection systems in ticks. The passive system detects the presence of CO_2_ and initiates active sensing by foreleg waving to detect potential hosts in the area. The foreleg sensory structures, including the HO, then seek out host emitted signals such as odors, body heat, and mechanosensory touch stimuli. Further studies on tick sensory physiology and neuronal innervation and projections are needed to unravel the definitive role of these two sensory pathways.

## Acknowledgments

I thank Wolfgang Stein (Illinois State University), Fernando Vonhoff (UMBC), and Kim-Ann Saal (UMG Göttingen) for helpful discussions and comments on the manuscript, Silvio Rizzoli (UMG Göttingen), Mark Frye (UCLA), and Wolfgang Stein for support and mentorship. Many thanks also to the Grass Foundation and 2021 Grass Fellows Bernardo Pinto, Duncan Leitch, Oscar Arenas Sabogal, Luis Bezares Calderon, as well as the 2021 Grass Director Melissa Coleman and Associate Director Laura Cocas and her postdoc Daniela Moura as well as the Trustees of the Foundation for support and constructive feedback. This work was supported by the Grass Foundation during the 2021 Grass Fellowship and by the UCLA Marion Bowen Postdoctoral Award.

## Author Contributions

C.S. designed experiments, collected data, analyzed data and wrote analysis codes, prepared figures, wrote the paper, and acquired funding.

## Data and Materials Availability

Supplemental Table S7 contains a list of all materials used. This paper does not report original code. Any additional information required to reanalyze the data reported in this paper is available from the lead contact upon request.

## MATERIAL AND METHODS

### Experimental Model Details

#### Tick purchasing and maintenance

Pathogen-free unfed virgin adult male and female *Ixodes scapularis* ticks were purchased from Oklahoma State University Centralized Tick Rearing Facility (Stillwater, OK, USA) approximately 6 weeks post-molting. Upon arrival, the animals were housed for 4 weeks under controlled laboratory conditions prior to experiments. Ticks were housed in 60 ml aseptic screw cap plastic vials (VWR, 216-1822P) with a mesh-covered 25 mm round hole in the lid. Each vial contained a maximum of 10 ticks. Males and females were maintained in separate vials. The tick vials were kept in transparent, airtight food storage containers (5.4 Liters, Emsa Clips & Close) at 99% relative humidity (saturated NaSO4 solution). Food containers containing the tick vials were kept inside an incubator (Binder, KBW E5.1) with a photoperiod of 14 hours light, 10 hours darkness, and circadian temperature cycling (21 °C, 6 am to 8 pm, lights on; 18 °C, 8 pm to 6 am, lights off). The animals were used for experiments 10 to 18 weeks after molting, never received blood, and were always separated by gender. The data presented were collected on ticks from 7 shipments over 18 months. We found no behavioral differences between ticks from different shipments, nor did we observe seasonal variation in CO_2_ responses of the lab-raised ticks used for this study.

### Experimental Setup

#### Multi-animal CO_*2*_ test assay

All experiments were performed during the day between 10 am and 4 pm at conditioned room temperature (21 ± 0.5 °C) and 40 to 60% relative humidity. For each experiment, five male or female *Ixodes scapularis* were placed on the test area (30 x 30 cm) inside a custom-made airtight Plexiglass box (Figure 1A, 40 x 40 x 30 cm) using a soft paintbrush without prior immobilization. Ticks were not pre-selected by, for example, blowing on them and seeing if they started to walk – a procedure occasionally used for selecting ticks for choice experiments. The walking surface of the test area was lined with white vinyl foil (VViViD XPO, matte finish) to ensure good contrast for video monitoring and a sufficient grip. The walking surface was cleaned with 70% ethanol before ticks were placed in the arena. A water moat prevented ticks from escaping the test area.

Ticks were allowed to acclimate for 30 minutes before the experiments. To test tick responses to different CO_2_ concentrations (Figures 1 & 2), we performed five experimental blocks on the same group of animals. We exposed ticks to an air puff as a negative control and to 1, 2, 4, and 8 % CO_2_. The treatment order was not randomized in these experiments. Ticks were first exposed to the air puff, followed by ascending CO_2_ treatments. CO_2_ was increased globally in the arena and there was no detectable gradient (see CO_2_ measurements). After acclimating, ticks were video monitored for 5 minutes in ambient air (2 frames/s) before the 5-minute-long treatment started. The animals were allowed to rest for 20 minutes between each experimental block, where the CO_2_ concentration was reset to ambient air levels (350-600 ppm) by opening the arena door. In experiments where treatments were only an air puff and 4% CO_2_ (Figures 3-5) stimuli were applied in randomized order.

#### Single-animal tick-on-a-stick assay

Experiments were carried out in the same Plexiglass box as the multi-animal experiments, with identical equipment. The test area was replaced with a floating ball setup, modified after Loesche and Reiser (2021). To facilitate handling, single male or female ticks were briefly immobilized through cold treatment. Animals were then tethered to a metal rod for fixation. In short, immobilized ticks were picked up with a paintbrush, placed into a movable cylindrical cavity (‘sarcophagus’) machined from solid brass, and positioned under a stereomicroscope (Zeiss, Stemi 2000). After aligning the tick in the sarcophagus, a thin metal rod was attached to the middle of the Scutum with a small drop of UV-curable glue (Bondic, UV liquid plastic). The tethered tick was then mounted on a micromanipulator (Siskiyou, MX10) for precise positioning along the three translational axes on top of an air-supported ball (General Plastics Manufacturing Company, Last-A-Foam FR-7120, 20 mm diameter) that served as an omnidirectional treadmill. The air supporting the ball was carbon-filtered with 350 to 600 ppm CO_2_. Care was taken that each tick was not cold immobilized for more than five minutes. Ticks were placed on the ball in a horizontal position.

Experiments started after 30 minutes of acclimatization at room temperature (21 ± 0.5 °C) and 40 to 60% relative humidity. We carried out two experimental blocks on the respective individual animals. Ticks were exposed to either an air puff or 4% CO_2_ in randomized order and video-monitored for 4 minutes (5 frames/s, 2 minutes before and after treatment). Only the last minute in the ambient air and the first minute during the treatment were used for the analysis (Figure 3).

### CO**_2_** measurements

To quickly change the CO_2_ concentration inside the arena and with as little gradient as possible, 100% CO_2_ was introduced into the Plexiglas box under high pressure (40 Liters/min). A diffusion pad (Genesee Scientific, #59-119) attached to one of the side walls of the box ensured broad CO_2_ distribution. The CO_2_ inflow was timed with a solenoid valve (Festo, VZWD Series) controlled by a timing relay (Panasonic, LT4H Series).

A flow meter (Vögtlin, Q-Flow 140 Series) was connected upstream of the solenoid valve to minimize disparity in CO_2_ concentrations between experiments. A computer fan with speed control (12 x 12 cm, Wathai) next to the diffusion pad provided continuous air movement and CO_2_ distribution. During the experiments the fan was constantly running on low speed (∼ 0.2 m/s airflow). This allowed sufficient airflow without creating wind and interfering with tick behavior.

CO_2_ was measured with an infrared CO_2_ Meter (Vaisala Inc., model GM70 + GMP251 Probe) at a sampling rate of 1 reading/second. To ensure that CO_2_ was evenly distributed in the arena and that the measured CO_2_ concentration corresponded to the concentration the ticks were experiencing, the CO_2_ meter was placed at 6 different locations in the arena (on the four side walls, the ceiling, and directly on the test area). For each CO_2_ sampling location three measurements were taken for each of the four CO_2_ concentrations tested and measurements were compared (data not shown). Irrespective of the CO_2_ concentration, measurements differed by ±50-250 ppm on average between sampling locations, which is within the measurement uncertainty of the CO_2_ probe (±0.1% / 1000 ppm at 5% CO_2_; ±0.2% / 2000 ppm at 8% CO_2_) and was therefore ignored. During the experiments, CO_2_ was measured at a mid-height at the rear side of the arena, with the probe placed slightly higher than the test area. CO_2_ was measured in ppm, but is reported as percentage values throughout the manuscript. CO_2_ delivery times were: 0.58 s for 10,000 ppm (1%), 1.25 s for 20,000 ppm (2%), 2.70 s for 40,000 ppm (4%), 5.75 s for 80,000 ppm (8%), each with a maximum deviation of ± 500 ppm.

### Air puff

We used a puff of carbon-filtered compressed air as a negative control to distinguish possible CO_2_-elicited behavioral changes from reactions to increased air movement during the onset of the treatments. The carbon-filtered air had the same pressure as the CO_2_ (40 Liters/min) and was introduced the same way into the Plexiglas box. Air puff duration was 5.75 s, the same time as for the 8% CO_2_ treatment. Before the air puff treatment, hoses were flushed extensively with air to remove CO_2_ residues. The CO_2_ concentration of the carbon-filtered compressed air did not differ from ambient air (350-600 ppm respectively).

### Video monitoring

It is questionable whether *Ixodes scapularis* has a rudimentary vision. As a precaution, all experiments were performed in darkness under red LED lighting (640-670nm, BestLEDStrip. com), a wavelength many arthropods cannot detect. Uniform lighting conditions were achieved by placing the LED lights around the test area. Ticks were video monitored with a monochrome camera (FLIR, Blackfly S) at 3.2 Megapixel (2,048 x 1,536 pixels). The camera was positioned outside on top of the Plexiglass box and animals were monitored from above. Multi-animal experiments were recorded with a 6mm/F1.4 fixed focal length lens (Edmund Optics, #67-709) at 2 frames/s. Single-animal tick-on-a-stick experiments were recorded with a 94mm macro lens (Infinity, #994100) at 5 frames/s.

### Classification of behavioral states

The following criteria were used to quantify behavioral changes in response to an air puff or CO_2_ treatment (Figures 2, 4F, and 5E):

#### Stationary

Animals that moved less than 5 mm within 2 minutes before the start of the treatments.

#### Horizontal questing

Stationary animals with raised forelegs for at least 1minute (example in Figure 1B).

#### Walking

Animals that moved more than 5 mm within 2 minutes before the start of the treatments.

#### Walking faster/slower

These behaviors only apply to animals that showed a change in walking speed during exposure to an air puff or CO_2_ compared to ambient air (= control condition). The mean walking speed (Δv_air_) and standard deviation (SD_air_) over 60 seconds immediately before treatment were calculated. Faster walking comprises animals whose mean walking speed during the treatment (Δv_treat_) was greater than the mean walking speed plus standard deviation in ambient air (Δv_treat_ > Δv_air_ + SD_air_). Slower walking comprises animals whose mean walking speed during the treatment was smaller than the mean walking speed minus standard deviation in ambient air (Δv_treat_ < Δv_air_ – SD_air_). Walking animals that did not meet either criterion were further classified as walking.

### Wax application and foreleg amputation

In some experiments, the foreleg tarsi (Figures 4, S4) and the mouth (Figure S6) were covered to test the contribution of the Haller’s and the palpal organ to CO_2_ detection. We used a paraffin-based casting wax (Kerzenkiste, #10815) with a low melting point of ∼50 °C. For a better visibility, the wax was dyed blue with liquid wax paint (Kerzenkiste, #10781-1-005). To minimize the risk of tissue damage from heat, the wax was pre-melted in a wax warmer at 55 °C (Amazon, Nivlan wax warmer). For application, the melted wax was picked up with a custom-made heating loop (diameter 0.6 mm, 0.19 mm silver wire) attached to a low-temperature soldering iron (Weller, WP80). This ensured that the wax did not harden between pickup and application. At least three layers were required to cover the long and wall-pored sensilla of the HO capsule completely. Cold immobilization and the brass sarcophagus used to glue ticks for the tick-on-a-stick experiments proved unhandy for wax application due to the condensation of atmospheric moisture on the cold surfaces and restricted access to the forelegs and mouthparts. It was thus inevitable to immobilize ticks with 100% CO_2_ using a diffusing pad (Genesee Scientific, #59-119) positioned under a stereomicroscope (Zeiss, Stemi 2000). Care was taken that ticks were not exposed to the CO_2_ for longer than 5 minutes. Ticks recovered within one to two minutes from the immobilization and were allowed to recover in the incubator at 21°C and 99% relative humidity for two hours before being tested. CO_2_ immobilization had no detectable influence on tick behavior. Ticks immobilized with CO_2_ for wax application on the foreleg patella (Supp. Figure S5) showed similar behavioral responses to 4% CO_2_ than not immobilized ticks (Figure 1).

In some experiments, the foreleg tarsi and HOs were amputated. Ticks were immobilized on CO_2_ for amputation as described above, and the tarsi were cut off with spring scissors (FST, Vannas Spring Scissors). After amputation, the animals were allowed to recover in the incubator at 21°C and 99% relative humidity for two hours before being tested.

### CO_**2**_ probe covering

To test whether the wax used to cover the foreleg HO and mouthparts is impermeable to CO_2_, we coated the CO_2_ probe with wax. We used one of the 60 ml tick housing plastic vials (VWR, 216-1822P) with a 25 mm mesh-covered hole in the lid. The lid was coated with a thin layer of the low-melting-point casting wax used to cover tick body parts. Therefore, the lid was briefly dipped into the melted wax in the wax warmer. A second hole was drilled into the bottom of the plastic vial where the CO_2_ probe was inserted. A clipped balloon (pink wrap in Supp. Figure S4D) and several layers of parafilm were used to seal the hole where the CO_2_ probe was inserted and the lid. The covered probe was then placed inside the Plexiglass behavior arena, and CO2 was added to the box for 2.7 and 6 seconds (3 trials each), corresponding to a CO_2_ concentration of 4% and 10%. CO_2_ concentration was reset to ambient air levels between trials by opening the arena door. The same experiment was carried out with a non-wax-covered lid to rule out that the CO_2_ measurements were affected by the attached plastic vial. A new lid was screwed onto the vial without affecting the balloon/parafilm seal.

### Data Analysis

#### Tracking

Reconstruction of the animals’ x-y body and foreleg positions was achieved with the Tracker Video Analysis and Modeling Tool (Open-Source Physics, Version 6.1.1). In multi-animal experiments the center of the body (center of mass) was tracked, in single-animal experiments the tip of the left and right forelegs. The tracked x-y coordinates were further processed and analyzed with custom-written MATLAB scripts (MathWorks, Version R2023a).

#### Heatmaps and Coefficient of Variation

Walking speed (mm/s) was calculated from x-y coordinates. Values were normalized to compare changes in walking speed across animals. For this, the data for the 10 minutes long experiments were binned (bin width = 30 frames / 15s) by calculating the mean velocity for each bin and each animal. Data from each animal were normalized to the respective mean of the experiment’s first 10 bins / 150 seconds.

The coefficient of variation (CV) was calculated from the normalized binned data using:

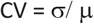

with *μ* as the mean of each bin and *σ* as the respective standard deviation. A CV unequal 1 means the data deviated from the bin’s mean value. Walking speed reductions lead to CVs < 1, and speed increases to CVs > 1.

#### Manual video analysis

To quantify the number of ticks that started to raise their forelegs or body, manual video analysis was necessary. The analysis was performed blindly by encrypting the file names and selecting them randomly (ImageJ, Blind Analysis Tool). Two minutes pre- and post-treatment of the 10-minute-long recordings were analyzed.

#### Foreleg positions

Foreleg angular positions were determined by placing a virtual polar coordinate system through the basal part of the animals’ mouthparts (base capitula, see Figure 3A). The average body length was used to calibrate the x-y coordinate system in the Tracker software (tip of the mouth to alloscutum; females 4 mm, males 3 mm). The four-quadrant inverse tangent in degrees was calculated from tracked x-y coordinates. 0° corresponds to forelegs exactly parallel to the midline. Negative angles are left and positive angles right. For x-y foreleg positions (Figure 3D & 3F), distances are given in mm. 0 mm corresponds to the axis intersection point at the basal part of the animals’ mouthparts.

#### Foreleg waving frequency

We used the position-time diagrams (Figure 3B & S3) to detect local maxima of angular positions of the left and right foreleg tips as a function of time. Minimum peak distance was 0.3 seconds. The waving frequency (Figure 3C) was calculated from the total number of local maxima over 60 seconds before and after the onset of the treatment.

#### Comparison of walking speed

To compare initial walking speeds, the mean walking speed over 5 minutes in ambient air was calculated for all animals that walked. In contrast, to compare walking speeds during the 4% CO_2_ treatment only walking and faster walking animals were included and the mean walking speed over 5 minutes was calculated.

#### Density distribution of the forelegs

The probability density estimate of the angular distribution of the forelegs (Figure 3E) was calculated and plotted with the Raincloud Plot extension for MATLAB (Allen et al., 2021). Total density of the foreleg tip x-y positions (Figure 3F) was determined by reducing the x-y coordinate system to one axis (x and y, respectively). The total length of the axis (8 mm, -4 mm to +4mm) was divided into 100 x 1 pixels (0.08 mm/ pixel). For each pixel, the sum of the x-y positions of the left and right foreleg tip was calculated (= total density). Data was not normalized and is given in mm distance from the axes intersection (the basal part of the animal’s mouthparts). The data were smoothed with a 5-pixel moving average filter.

### Quantification and Statistical Analysis

Statistical analysis was performed with MATLAB (MathWorks, Version R2023a). The Shapiro–Wilk test was used to test data for normal distribution. In the text, we report mean values ± standard deviation. All data tested for statistical differences were normally distributed. Results from t-tests are reported as t (degrees of freedom) = t value, p value either in the text or in the figure legends. Final figures were prepared with Adobe Illustrator (version 24.2.3).

