## Supplementary Material for "The black-legged tick *Ixodes scapularis* detects CO_2_ without the Haller’s organ"

<u>*This PDF file includes:*</u>

*Figs. S1 to S6*

*Table S7*

*Captions for supplemental videos S7 to S8*

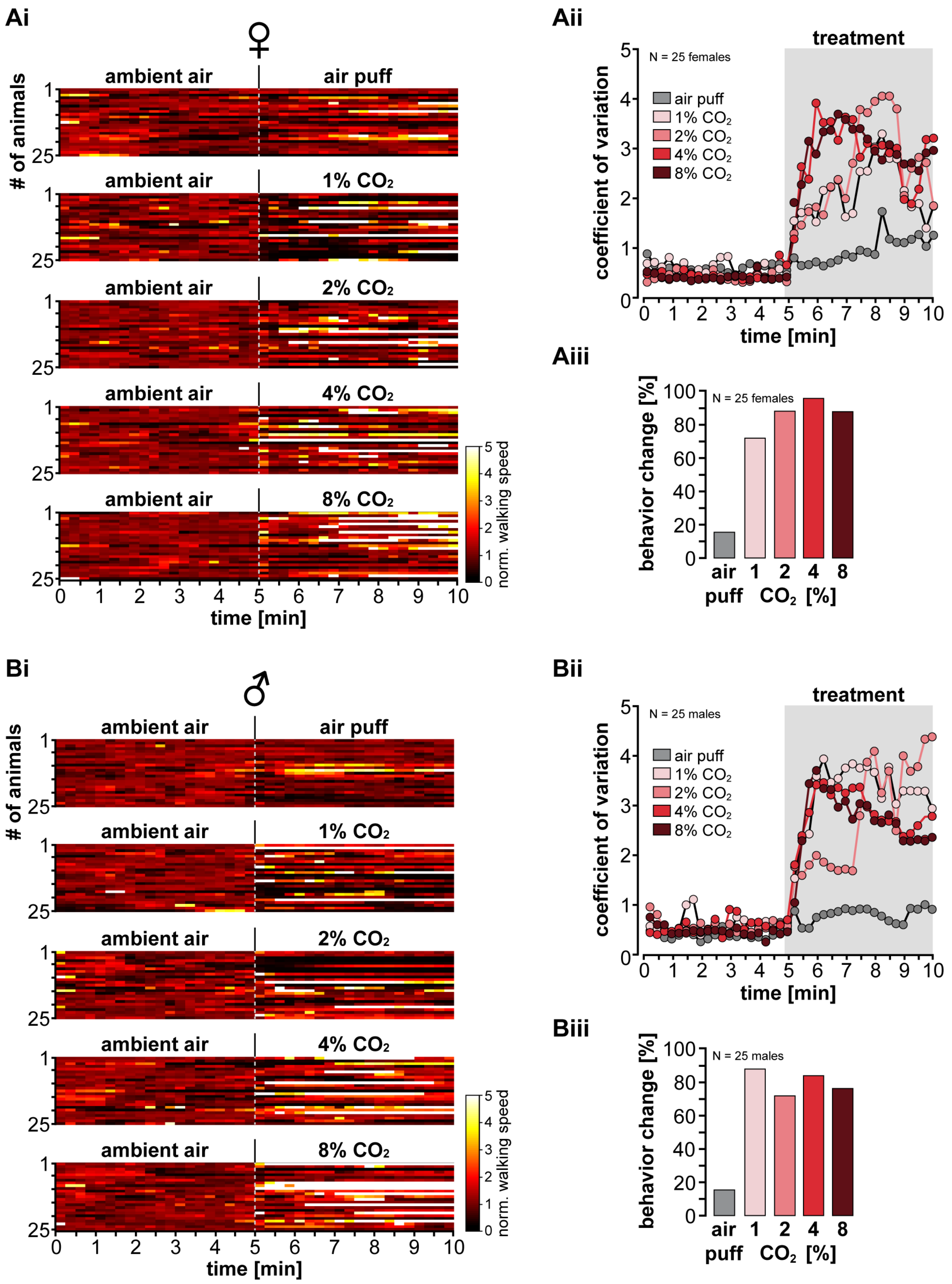


**Figure S1 (Figure 1 supplement):** ***Ixodes scapularis*** **reaction to $CO_2$ separated by gender.**

(**A**) (i) Analysis of changes in normalized walking speed before (ambient air) and during exposure to different $CO_2$ concentrations/air puff for the 25 females tested. Brighte r colors represent faster walking speeds. Animals were first monitored in ambient air (=control condition) before the treatment started (1, 2, 4, 8% $CO_2$ or an air puff as negative control). The dashed gray line depicts the beginning of the treatment. Bin width = 30 frames, sampling rate = 2 frames/s. Data from each animal were normalized to the respective mean of bins 1 to 10. (ii) Coefficient of variation for the data presented in (i). Dots represent mean values. The gray rectangle illustrates the start and duration of the treatment. (iii) Manual video analysis of behavioral changes of the same 25 females.

(**B**) Same as panel A but for the 25 male *I. scapularis* tested.

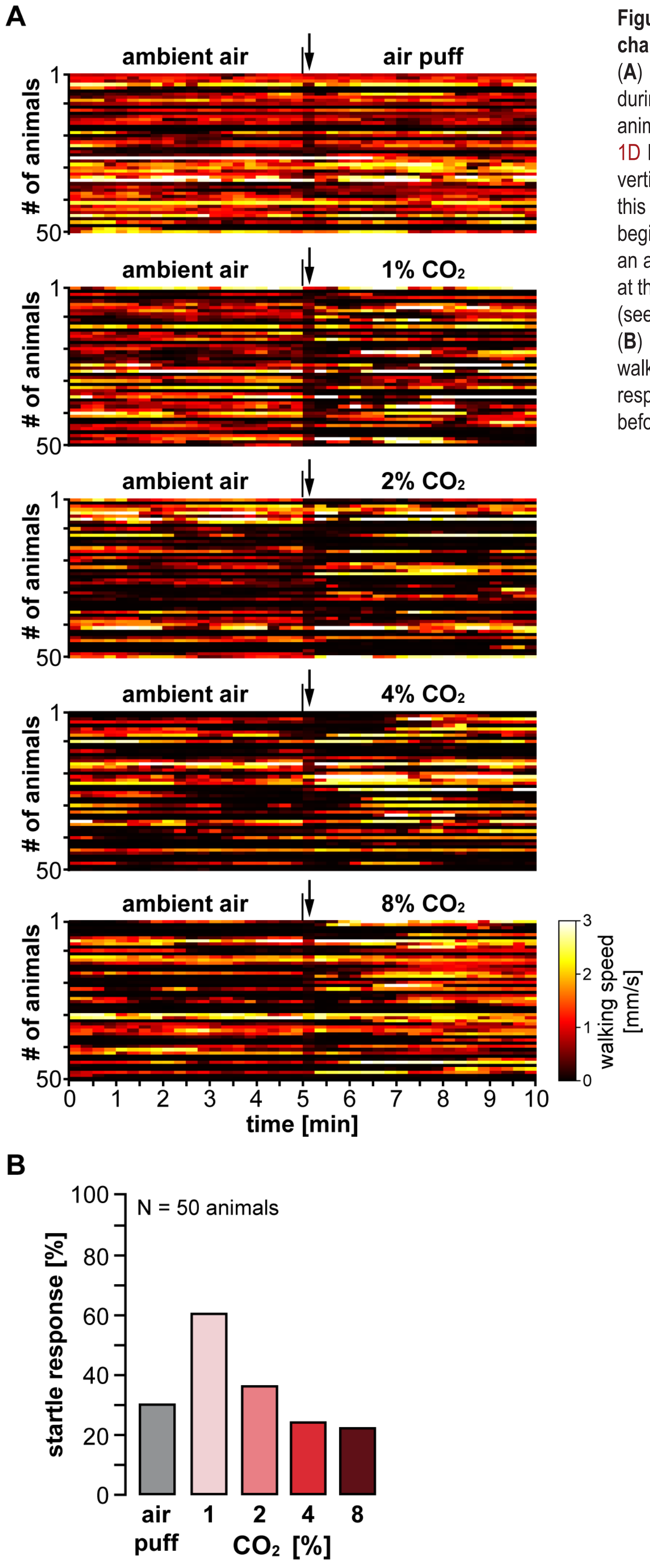


**Figure S2 (Figure 1 supplement): *Ixodes scapularis* responds to changes in airflow**

(**A**) Analysis of changes in walking speed before (ambient air) and during exposure to different $CO_2$ concentrations/air puff for the 50 animals tested (25 males and 25 females). Same data as in Figure 1D but not normalized. Brighter colors represent faster walking. The vertical dashed line, as seen in other heatmaps, was removed from this plot for better visualization. Here the vertical black lines depict the beginning of the treatment. Regardless of whether the treatment was an air puff or $CO_2$, walking speed decreased briefly (1 bin / 15 s) right at the onset of the stimulus – recognizable by the darker-colored band (see arrow).

(**B**) Bar graph depicting the percentage of animals that reduced their walking speed by half or more right at the start of treatment (= startle response). The mean walking speeds over 15 seconds immediately before and after the start of treatment were compared.

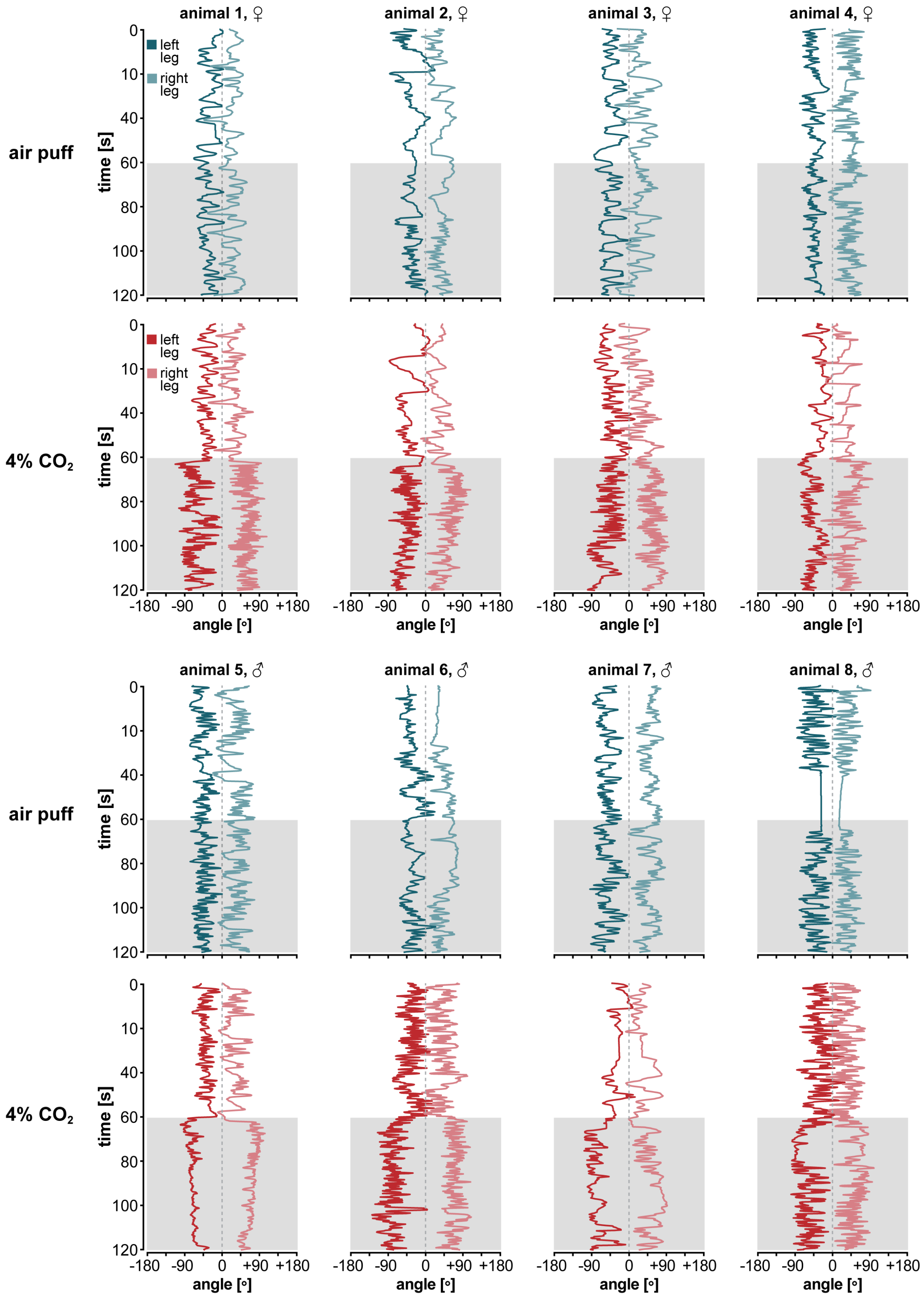


**Figure S3 (Figure 3 supplement): $CO_2$ elicits foreleg waving.**
Space-time diagrams depicting the angular positions of the left and right foreleg tip as a function of time for the 8 animals tested (row 1-2: females, row 3-4: males). Leg movement was monitored over 60 seconds in ambient air (white background) before the treatment started (gray rectangle, 60 seconds; air puff, blue, 4% $CO_2$, red). Sampling rate = 10 frames/s. See Figure 3A in the main article for polar coordinate system details.

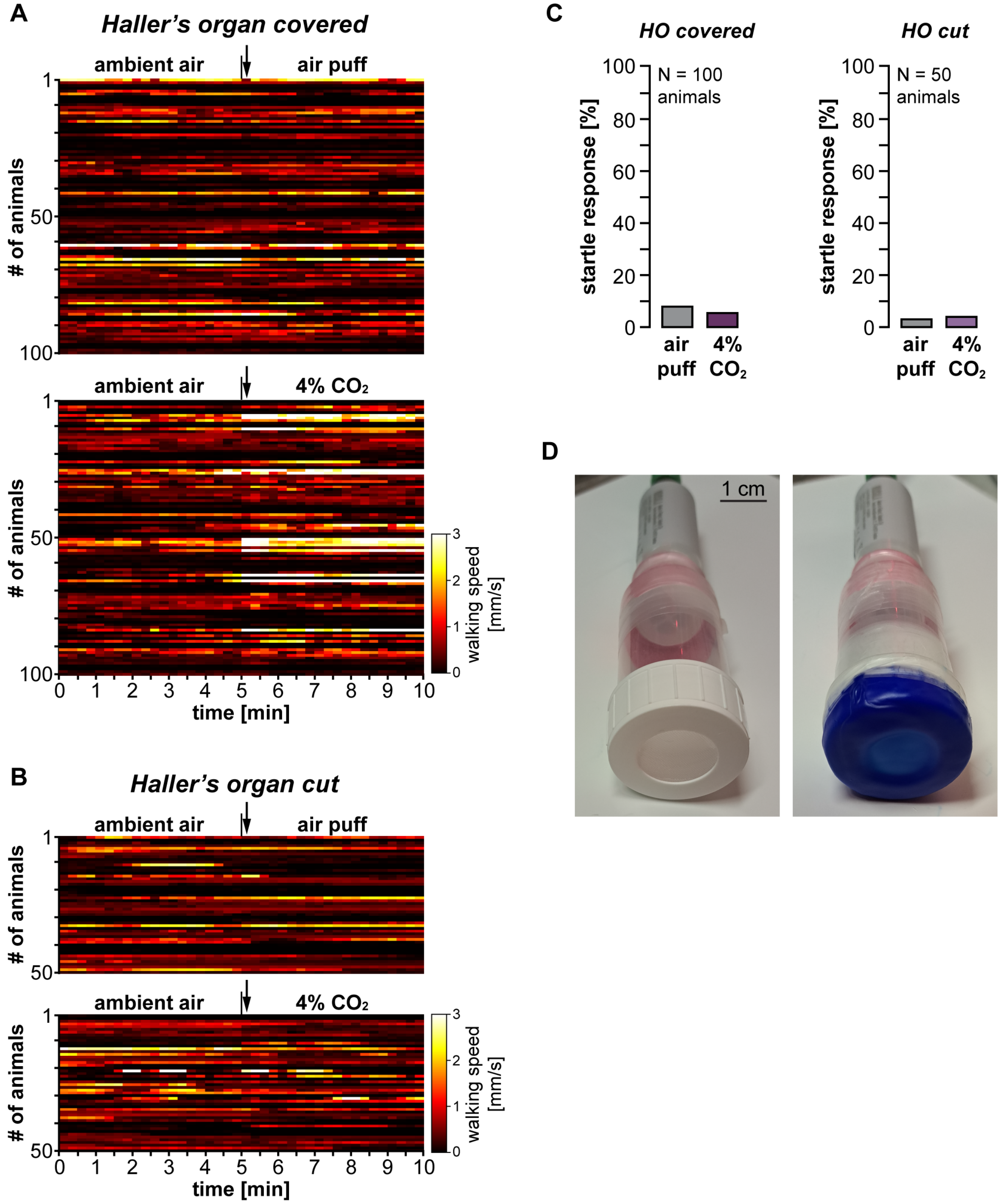


**Figure S4 (Figure 4 & 5 supplement): Disabling the Haller's organ suppresses startle responses.**

(**A**) Analysis of changes in walking speed before (ambient air) and during exposure to different $CO_2$ concentrations/air puff for the 100 animals tested with wax-covered forelegs (50 males and 50 females). Same data as in Figure 4C but not normalized. Brighter colors represent faster walking. The vertical black lines depict the beginning of the treatment. With covered forelegs, walking speed no longer decreased briefly at the onset of the treatment (see arrow) - the darker-colored band is no longer present. See Supplemental Figure S2 for comparison.

(**B**) Analysis of changes in walking speed for the 50 animals tested with cut forelegs (25 males and 25 females). Same data as in Figure 5C but not normalized. With cut forelegs, *I. scapularis* no longer decrease their walking speed at the onset of the treatment, and the darker-colored band is no longer present.

(**C**) Bar graphs depicting the percentage of animals that reduced their walking speed by half or more right at the start of treatment (= startle response). Left: animals with wax-covered Haller's organ. Right: animals with transected Haller's organ. The mean walking speeds over 15 seconds immediately before and after the start of treatment were compared.

(**D**) Pictures of the non-wax-covered $CO_2$ probe (left) and the wax-covered $CO_2$ probe, as used for the control experiments presented in Figure 4F.

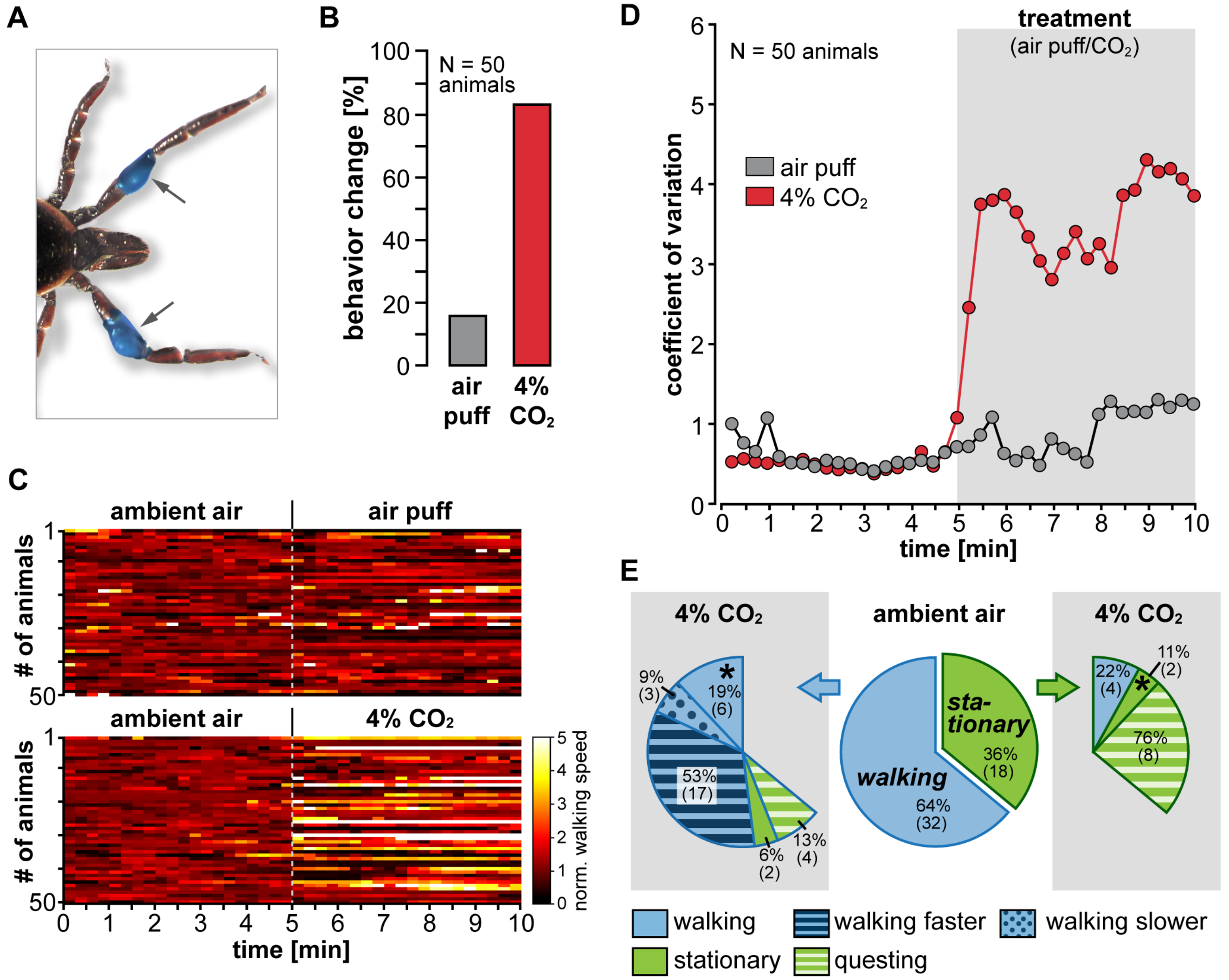


**Figure S5 (Figure 4 supplement): Applying wax does not affect behavior.**

(**A**) Photo showing the position of the blue-colored low melting point wax on the foreleg patella. Approximately the same amount of wax was applied as in experiments covering the tarsi.

(**B**) Manual video analysis of behavioral changes for 50 adult *I. scapularis* (25 males and 25 females) with wax added to the patella. Behavioral changes include changes in walking speed and foreleg/body position.

(**C**) Heatmap showing the normalized walking speed. Brighter colors represent faster walking. Data from each animal were normalized to the respective mean of bins 1 to 10.

(**D**) Coefficient of variation for the data presented in panel D. Dots represent mean values. The gray rectangle illustrates the start and duration of the treatment. Animals with wax added to the patella did show similar responses to the 4% $CO_2$ treatment as animals with wax-free forelegs. See Figure 1 for comparison.

(**E**) Pie charts showing the percentage of exhibited behaviors for 50 animals with wax on the patella before (ambient air) and during exposure to 4% $CO_2$. Animals were sorted based on their dominant behavior in ambient air (walking or stationary).

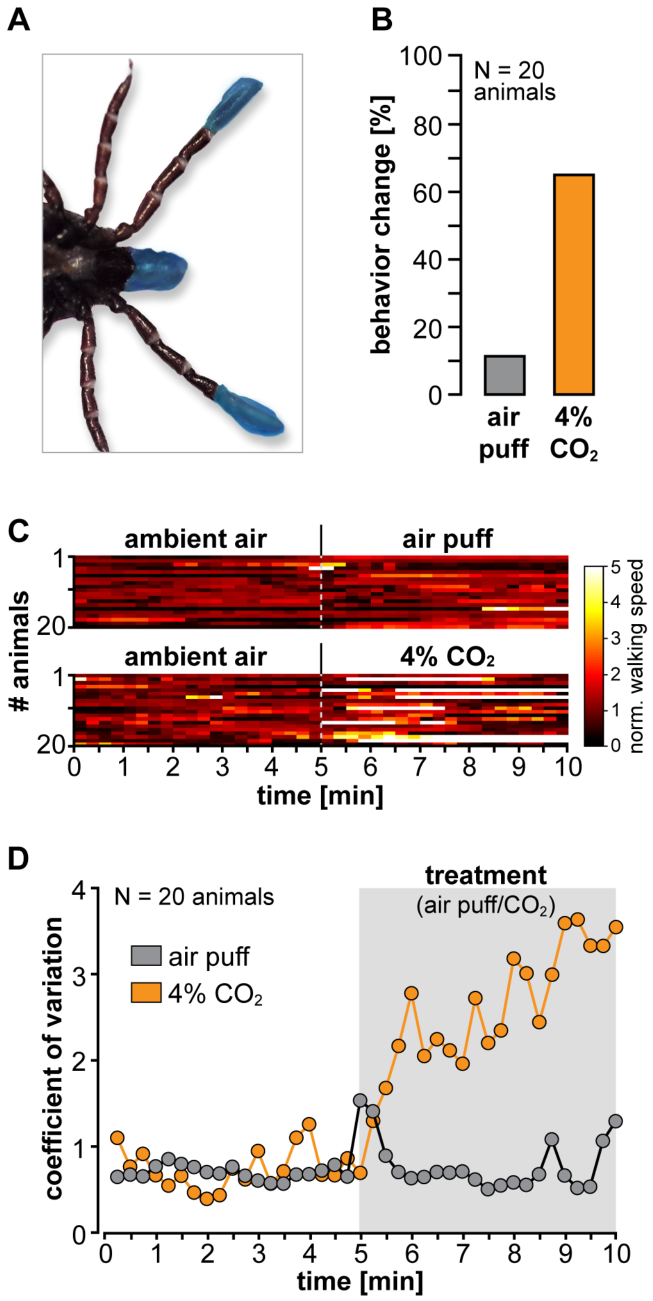


**Figure S6 (Figure 4 & 5 supplements): Disabling the palp sensilla does not eliminate $CO_2$ responses.**
(**A**) Photo showing an adult *I. scapularis* female (ventral view) with wax-covered mouthparts (palpal organ) and foreleg tarsi (Haller's organ).
(**B**) Manual video analysis of behavioral changes for 20 adult *I. scapularis* (10 males and 10 females) with wax-covered palpal and Haller's organ. Behavioral changes include changes in walking speed and foreleg/body position.
(**C**) Heatmap showing the normalized walking speed for the same 20 animals as in panel B in ambient air and during 4% $CO_2$ exposure. Brighter colors represent faster walking. Data from each animal were normalized to the respective mean of bins 1 to 10.
(**D**) Coefficient of variation for the data presented in panel D. Dots represent mean values. The gray rectangle illustrates the start of the treatment. Animals with wax-covered palpal and Haller's organ did show similar responses to the 4% $CO_2$ treatment as wax-free animals. See Figure 1 for comparison.

**Table S7: Resources Table, listing the reagents and materials used in the study**

| *REAGENT or RESOURCE* | *SOURCE* | *IDENTIFIER* |
|---|---|---|
| ***Experimental organisms*** | | |
| *Ixodes scapularis*, male and female unfed adults | Oklahoma State University Centralized Tick Rearing Facility (Stillwater, OK, USA) | n/A |
| ***Software and algorithms*** | | |
| MATLAB R2023a | MathWorks | |
| Raincloud Plot extension for MATLAB | Allen, M. et al. (2019) | doi:10.12688/wellcomeopenres.15191.1 |
| Tracker Video Analysis and Modeling Tool, Version 6.1.1 | Open-Source Physics | https://physlets.org/tracker/ |
| ImageJ, Blind Analysis Tool Plugin | GitHub | https://imagej.net/plugins/blind-analysis-tools |
| Adobe Illustrator 2023 | Adobe | |
| ***Other*** | | |
| Bondic UV liquid plastic glue | Amazon | ASIN B00QU5M4MG |
| computer fan with speed control, 12 x 12 cm, Wathai | Amazon | ASIN B07VYGQPCZ |
| Nivlan wax warmer | Amazon | ASIN B082KDNK3Z |
| 60 ml aseptic screw cap plastic vials | avantor / VWR | Cat# 216-1822P |
| red LED lighting strip, 640-670nm | BestLEDStrip | https://www.bestledstrip.com/12v-red-led-strip-light-5050-60-5m/ |
| Incubator/Climatic chamber | Binder GmbH | Cat# KBW E5.1 |
| stereomicroscope | Carl Zeiss AG | Stemi 2000 |
| Panasonic timing relay | Conrad | Cat# 505207 |
| Handheld $CO_2$ Meter, Vaisala Inc. | Driesen + Kern GmbH | Cat# MI70 |
| infrared $CO_2$ probe 0-100.000 ppm, Vaisala Inc. | Driesen + Kern GmbH | GMP251B1C0B0N1 |
| 6mm/F1.4 fixed focal length lens | Edmund Optics | Cat# 67-709 |
| 5.4 Liter airtight food storage containers, Clip & Close | Emsa | https://www.emsa.com/produkt/clip-close-frischhaltedosen-rechteckig |
| solenoid valve, VZWD Series | Festo | Cat# VZWD-L-M22C-M-G14-25-V-1P4-8 |
| Vannas-Tübingen Spring Scissors | Fine Science Tools | Cat# 15003-08 |
| monochrome computer vision camera, Blackfly S | FLIR | Cat# BFS-U3-32S4M |
| Air-supported ball | General Plastics Manufacturing Company | Last-A-Foam FR-7120 |
| Flystuff $CO_2$ Anesthetizing pad, 10.1 x 14 cm | Genesee Scientific | Cat# 59-119 |

| | | |
|---|---|---|
| 94mm macro lens InfiniStix | Infinity | Cat# 994100 |
| liquid wax paint, blue, 50 ml | Kerzenkiste | Cat# #10781-1-005 |
| paraffin-based casting wax | Kerzenkiste | Cat# 10815 |
| Three-axis micromanipulator | Siskiyou | MX10 |
| flow meter, Q-Flow 140 Serie | Vögtlin Instruments GmbH | Cat# 134-1342 |

**Video S8: Behavioral reactions to $CO_2$ for adult *Ixodes scapularis* with intact and disabled Haller's organ**

Shown are behavioral responses of 1) ticks with intact Haller's organ, 2) wax-covered forelegs, and 3) amputated forelegs. In each case, the reactions of 2 experiments are shown side by side for both walking and stationary ticks. Different ticks are shown in the individual video clips. The recordings with intact Haller's organ were made in a reduced experimental setup to obtain better video resolution for demonstration purposes. The two shown animals with intact Haller's organ are not included in the manuscript's data analysis. In contrast, the video clips with wax-covered and amputated Haller's organ are from actual experiments and taken from a greater distance resulting in lower resolution.

**Video S9: Example video of the single animal tick-on-a-stick CO2 responses**

Responses to ambient air and 4% $CO_2$ of an adult female *Ixodes scapularis*, tethered to a metal rod and placed on an air-supported ball that acts as an omnidirectional treadmill.
